# Genomic data and multi-species demographic modelling uncover past hybridization between currently allopatric freshwater species

**DOI:** 10.1101/585687

**Authors:** Sofia L. Mendes, Miguel P. Machado, Maria M. Coelho, Vitor C. Sousa

## Abstract

Evidence for ancient interspecific gene flow through hybridization has been reported in many animal and plant taxa based on genetic markers. The study of genomic patterns of closely related species with allopatric distributions allow to assess the relative importance of vicariant isolating events and past gene flow. Here, we investigated the role of gene flow in the evolutionary history of four closely related freshwater fish species with currently allopatric distributions in western Iberian rivers - *Squalius carolitertii, S. pyrenaicus, S. torgalensis* and *S. aradensis* - using a population genomics dataset of 23 562 SNPs from 48 individuals, obtained through genotyping by sequencing (GBS). We uncovered a species tree with two well differentiated clades: (i) *S. carolitertii* and *S. pyrenaicus*; and (ii) *S. torgalensis* and *S. aradensis*. By using D-statistics and demographic modelling based on the site frequency spectrum, comparing alternative demographic scenarios of hybrid origin, secondary contact and isolation, we found that the *S. pyrenaicus* North lineage is likely the result of an ancient hybridization event between *S. carolitertii* (contributing ~84%) and *S. pyrenaicus* South lineage (contributing ~16%), consistent with a hybrid speciation scenario. Furthermore, in the hybrid lineage we identify outlier loci potentially affected by selection favouring genes from each parental lineage at different genomic regions. Our results suggest that ancient hybridization can affect speciation and that freshwater fish species currently in allopatry are useful to study these processes.

## Introduction

How populations diverge and ultimately originate new species has always intrigued evolutionary biologists. Speciation is assumed to occur due to a continuous reduction in gene flow until reproductive isolation is achieved and populations maintain phenotypic and genetic distinctiveness (Coyne and Orr, 2004). This process might occur between geographically isolated populations in a strictly allopatric scenario, or in sympatry driven by ecological factors (Bush, 1975; Schluter, 2009). In either case (and the continuum in-between), genetic differences are expected to accumulate so that reproduction between the initial populations is no longer possible (Coyne and Orr, 2004).

Understanding the role of gene flow is key to explain how biological diversity is generated and species are formed. Historically, gene flow was considered a homogenizing force, mixing gene pools and counteracting the accumulation of divergence between populations. However, based on population genetics data, evidence for divergence in the presence of gene flow has accumulated (reviewed in Feder *et al*., 2012). For example, ecologically driven divergent selection has been shown to promote population divergence in the absence of barriers to gene flow in several animal taxa, including insects, fish and mammals (Linn *et al*., 2003; Gagnaire *et al*., 2013; Pfeifer *et al*., 2018).

While the role of hybridization between species in evolution has been discussed for quite some time (e.g. Anderson and Stebbins, 1954; Stebbins, 1959; Grant, 1981), advances in our ability to generate population genomics data and perform fine-scale analysis (Davey *et al*., 2011; da Fonseca *et al*., 2016; Payseur and Rieseberg, 2016) have shown that hybridization is more widespread than previously thought, especially in the animal kingdom (reviewed in Taylor and Larson, 2019). Hybrid individuals might be unfit in their parentals’ habitat or have reduced viability and fertility due to hybrid incompatibilities (Bateson-Dobzhansky-Muller incompatibilities (Dobzhansky, 1937; Muller, 1942)). However, mixing old alleles into new combinations through hybridization can fuel adaptation and speciation (e.g. Rieseberg *et al*., 2003; Hermansen *et al*., 2014; Wallbank *et al*., 2016; Richards and Martin, 2017; Svardal *et al*., 2020; reviewed in Marques *et al*., 2019). In this context, it is crucial to determine the timing and mode of gene flow. If hybridization between two species occurs at the time of origin of a third new reproductively isolated lineage, it is likely a case of hybrid speciation (reviewed in Mallet, 2007; Abbott *et al*., 2013; Vallejo-Marín and Hiscock, 2016). Instead, if hybridization occurs without producing a new lineage but results in introgression of genetic material through backcrossing, it is a case of secondary contact.

Due to their outstanding diversification and remarkable adaptive radiations, freshwater fish have been widely used as model systems to study speciation (e.g. reviewed in Bernardi, 2013 and Seehausen and Wagner, 2014). Different events fuelled this diversity, including transitions from marine to freshwater habitats (e.g. Jones et al. 2012; Terekhanova et al. 2014), adaptation to extreme environments (e.g. Pfenninger et al. 2015), differentiation along water depth clines (e.g. Barluenga et al. 2006; Gagnaire et al. 2013) and changes in the configuration of rivers and lakes over geological time (e.g. Sousa-Santos *et al*., 2019). Freshwater fish also stand out as a vertebrate group with high hybridization rates (Hubbs, 1955; Wallis *et al*., 2017). For example, hybridization events that promoted rapid adaptive radiations within lake systems have been documented (e.g. Meier *et al*., 2017; Svardal *et al*., 2020), as well as several instances of hybridization between native and invasive species (e.g. Nolte *et al*., 2006; Meraner *et al*., 2013).

In this work, we analyse the role of gene flow during the speciation process of four species of Iberian endemic chubs found in Western Iberian rivers: *Squalius carolitertii, Squalius pyrenaicus, Squalius torgalensis* and *Squalius aradensis* (Figure 1). As obligatory freshwater fish, their evolutionary history is intertwined with the geomorphological rearrangements of the river systems they inhabit. These species are distributed along a latitudinal environmental cline with increasing temperatures and propensity for drought from north to south, reflecting the transition from an Atlantic to a Mediterranean climate type (Gasith and Resh, 1999; Jesus *et al*., 2017). Two of them have rather wide distribution ranges: *Squalius carolitertii* (Doadrio, 1988) inhabits northern rivers down to the Mondego basin, while *Squalius pyrenaicus* (Gunther, 1868) has a more southern distribution (Tagus, Sado, Guadiana and south-eastern basins) (Coelho *et al*., 1995, 1998). Contrastingly, the two other species are confined to small basins in the southwest: *Squalius torgalensis* (Coelho *et al*., 1998) inhabits the Mira basin and *Squalius aradensis* (Coelho *et al*., 1998) basins in the extreme southwestern area (e.g. Arade) (Coelho *et al*., 1998). These two last species are “Critically Endangered” and *S. pyrenaicus* is “Endangered” in the Portuguese Red List (Cabral *et al*., 2005) and “Vulnerable” in the Spanish Red List (Doadrio, 2002).

**Figure 1.**
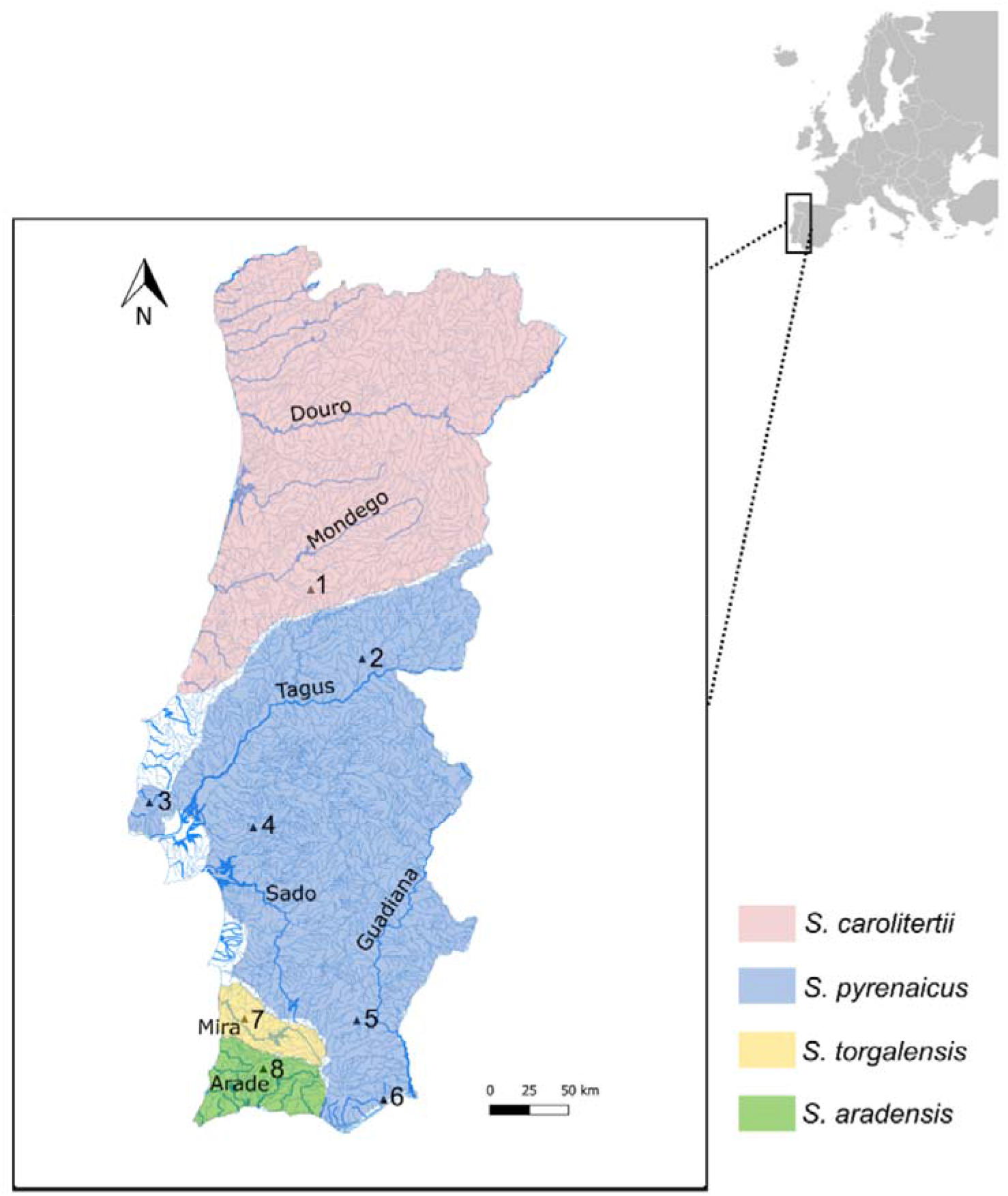
Distribution range of the four *Squalius* species in Portuguese rivers and sampling locations: (1) Mondego; (2) Ocreza; (3) Lizandro; (4) Canha; (5) Guadiana; (6) Almargem; (7) Mira; (8) Arade. Triangles correspond to the GPS coordinates of the sampling locations.

Although these species have allopatric distributions (Figure 1) associated with different river basins, studies based on mitochondrial and nuclear markers found phylogenetic incongruences between the two types of markers, which have been interpreted as possible past hybridization but remain unresolved (Brito *et al*., 1997; Waap *et al*., 2011; Sousa-Santos *et al*., 2019; Perea *et al*., 2020). Thus, this work had three major goals: (i) to characterize the genome-wide patterns of genetic differentiation and reconstruct the species tree for these four *Squalius* species; (ii) to investigate the role of gene flow in the evolution of these species, particularly to assess if previously reported incongruent mitochondrial and nuclear phylogenies can be explained by incomplete lineage sorting; and (iii) to perform demographic modelling to compare alternative scenarios, and to date and quantify past gene flow. To achieve these goals, we obtained genome-wide single nucleotide polymorphisms (SNPs) across the four species through Genotyping by Sequencing (GBS) and applied methods to quantify genetic differentiation, reconstruct the evolutionary relationship between species and test for evidence of gene flow. Furthermore, we performed demographic modelling to compare alternative models of isolation, hybrid origin and secondary contact, accounting for drift and shared ancestry (i.e., incomplete lineage sorting). We found support for a potential case of hybrid speciation in *S. pyrenaicus*, uncovering genomic regions that might have been under selection in the hybrid.

## Methods

### Sampling and sequencing

We sampled 65 individuals from 8 different locations, as displayed in Figure 1, including at least one sampling location from a representative river basin per species. *S. carolitertii* individuals were collected from the Mondego basin (n=10). In the northern part of *S. pyrenaicus* distribution individuals were collected from the Ocreza river (n=10) and Canha stream (n=10), both tributaries of the Tagus basin. Specimens were also collected in the Lizandro basin (n=10). From here on, we use “*S. pyrenaicus* North” to refer to *S. pyrenaicus* from Ocreza, Canha and Lizandro. In the southern part of the distribution, *S. pyrenaicus* was sampled in the Guadiana (n=2) and Almargem (n=8) basins, which we refer to as “*S. pyrenaicus* South”. *S. torgalensis* individuals were collected in the Mira basin (n=10) and *S. aradensis* individuals from the Arade basin (n=5). GPS coordinates of the sampling locations and fishing licenses from the Portuguese authority for conservation of endangered species [ICNF - Instituto de Conservação da Natureza e das Florestas] can be found in Table S1.

Fish were collected by electrofishing (300V, 4A) and total genomic DNA was extracted from fin clips using a phenol-chloroform protocol adapted from Taggart *et al*., 1992 and quantified using Qubit^®^ 2.0 Fluorometer (Live Technologies). Samples were subjected to a paired-end Genotyping by Sequencing (GBS) protocol (adapted from Elshire *et al*., 2011), performed at Beijing Genomics Institute (BGI, www.bgi.com). DNA was sent to the facility mixed with DNAstable Plus (Biomatrica) to preserve it at room temperature during shipment. Briefly, upon arrival, DNA was fragmented using the restriction enzyme ApeKI and the fragments were amplified after adaptor ligation (Elshire *et al*., 2011). The resulting library was sequenced using Illumina Hiseq2000.

### Multispecies SNP dataset

We assessed the quality of the sequences using FastQC (https://www.bioinformatics.babraham.ac.uk/projects/fastqc/) and MultiQC (Ewels *et al*., 2016). We then used *process_radtags* from Stacks v2.2 (Rochette *et al*., 2019) to trim all reads to 82 base pairs and discard reads with uncalled bases and low quality scores, using the default settings for window size (0.15x read length) and base quality threshold (10 Phred score). Given the absence of a reference genome for any of the species in this study, we built a catalog of all loci using the *denovo* approach from Stacks v2.2 (Rochette *et al*., 2019). We first tested different M and n parameter (see definition below) values for catalog construction using a subset of representative individuals (Figure S1), following the approach recommended by Paris et al. 2017. We based our parameter selection on accounting for differences between four species with an estimated divergence time of 14Mya (Sousa-Santos *et al*., 2019) and low genetic diversity (Almada and Sousa-Santos, 2010). Using all individuals for catalog construction, we required a minimum depth of coverage of 4x for every stack (set of exactly matching reads) (m=4) and allowed a maximum of 2 mismatches between initially formed stacks to be considered putative alleles of the same locus within each individual (M=2). We then allowed a maximum of 4 differences between stacks from different individuals (n=4) for them to be considered one locus on the catalog. Given the possibility that forward and reverse sequences of the same fragment were treated as different loci, similar reads within the catalog were clustered using CD-HIT-EST from the CD-HIT v4.7 package (Li and Godzik, 2006; Fu *et al*., 2012) with a word length of 6 and a sequence identity threshold of 0.85. The reads from each individual were aligned against the catalog using BWA-MEM from BWA v0.7.17-r1188 (Li, 2013) with default parameters. We sorted the output alignments and removed unmapped reads using Samtools v1.10 (Li and Durbin, 2009) and evaluated the alignments for each individual using Picard v2.18.13 (http://broadinstitute.github.io/picard/) (Table S2). To call genotypes for each individual at each site and identify SNPs we used the method implemented in Freebayes v1.2.0 (Garrison and Marth, 2012), discarding reads and bases with low quality and without using Hardy-Weinberg equilibrium priors (−p 2 --min-mapping-quality 30 --min-base-quality 20 --hwe-priors-off).

To discard sites and genotypes that are likely due to sequencing or mapping errors and to maximize the number of individuals with data, we applied a series of filters using a combination of options from VCFtools v0.1.15 (Danecek *et al*., 2011) and BCFtools v1.6 (Li *et al*., 2009). First, we kept only SNPs present in all sampling sites in at least 50% of the individuals. Second, genotypes with a depth of coverage (DP) outside of ¼ to 4 times the individual median DP and SNPs with excess of heterozygotes when pooling all individuals were removed. Third, individuals with more than 50% missing data were removed. Finally, only SNPs with a minor allele frequency (MAF) larger than 0.01 were kept (MAF≥0.01).

### Global patterns of genetic differentiation

To quantify the levels of differentiation between sampling locations, we calculated the pairwise F_ST_ using the Hudson estimator (Hudson *et al*., 1992). We investigated fine population structure using individual based methods: principal component analysis (PCA) and ADMIXTURE v1.3 (Alexander *et al*., 2009). PCA was performed using package LEA (Frichot and François, 2015) in RStudio v1.1.383 and R v3.4.4. We used ADMIXTURE to determine the ancestry proportion of each individual from a specified number of clusters (K), testing values of K between 1 and 8, performing 100 independent runs for each value of K. We identified the best value of K as the one with the lowest 5-fold cross-validation error. For 2≤K≤5, we assessed similarity across the 100 replicates using the Greedy algorithm implemented in CLUMPP v1.1.2 (Jakobsson and Rosenberg, 2007).

### Relationship between species and populations

We reconstructed a graph describing the relationships between the populations using TreeMix v1.13 (Pickrell and Pritchard, 2012). We explored models with no migration and up to two migration events. Since there is only one individual from *S. pyrenaicus* Guadiana in the final dataset (see Results), *S. pyrenaicus* South is represented only by *S. pyrenaicus* Almargem. The position of the root was not specified, thus the resulting trees are unrooted.

We also constructed a maximum likelihood phylogeny of the individuals using IQ-TREE v1.6.12 (Nguyen *et al*., 2015). We used vcf2phylip v2.0 (Ortiz, 2019) to convert the VCF file into PHYLIP format. We used ModelFinder (Kalyaanamoorthy *et al*., 2017) implemented on IQ-TREE to determine the best substitution model, limiting the search to models with ascertainment bias correction (+ASC). According to the corrected Akaike Information Criterion (AIC), the best model was GTR+F+ASC+R2, which was used to construct the phylogeny with 5 000 standard non-parametric bootstrap replicates. The resulting best tree was visualized with midpoint-rooting in FigTree v1.4.4 (http://tree.bio.ed.ac.uk/software/figtree/).

### Effect of linked SNPs

To verify if the results were influenced by potential linkage between SNPs, we produced a dataset minimizing the number of linked SNPs by dividing the catalog into blocks of 200 base pairs, which is larger than the mean size of GBS loci, and sampling one SNP per block. We selected the SNP with less missing data per block. Using this “single SNP” dataset, we repeated analyses that might be affected when SNPs are not independent (PCA, ADMIXTURE and TreeMix).

### Detection of introgression between *S. carolitertii* and *S. pyrenaicus*

To test for past introgression between *S. carolitertii* and *S. pyrenaicus* we used the D-statistic (Durand *et al*., 2011) with four different combinations of populations, using both *S. aradensis* and *S. torgalensis* as outgroup in all cases. First, we tested either *S. carolitertii* or S. *pyrenaicus* South as the possible sources of introgression into *S. pyrenaicus* North. To test for geographical cline in admixture proportions in *S. pyrenaicus* North, we tested if the northernmost sampling site of *S. pyrenaicus* (Ocreza - Figure 1) showed more shared alleles with *S. carolitertii* than the other *S. pyrenaicus* North (Lizandro and Canha). Finally, we also considered all *S. pyrenaicus* North as sister populations and *S. carolitertii* as the potential source of introgressed genes.

As in the TreeMix analysis, we used *S. pyrenaicus* Almargem as the *S. pyrenaicus* South population. Significance of D-statistic values was assessed using a block jackknife approach (Soraggi *et al*., 2018), dividing the dataset into 25 blocks with a similar number of SNPs. These computations were performed in RStudio v1.1.383 and R v3.4.4 using custom scripts.

If introgression between populations occurred in the relatively recent past, we would expect individuals within the same population to show different degrees of introgression. To test this hypothesis, we calculated the D-statistic for each individual of *S. pyrenaicus* North for the same scenarios as above.

### Demographic modelling of the divergence of *S. carolitertii* and *S. pyrenaicus*

We compared alternative divergence scenarios of *S. pyrenaicus* and *S. carolitertii* using the composite likelihood method based on the site frequency spectrum (SFS) implemented in *fastsimcoal2* v2.6 (Excoffier *et al*., 2013).

First, we compared the fit of three models with and without admixture to the observed SFS (see Figure S2 for parameters inferred in each model). We compared a model that assumes *S. pyrenaicus* North received a contribution alpha (α) from *S. pyrenaicus* South and 1-alpha (1-α) from *S. carolitertii* at the time of the split (Figure S2A) with models without admixture (Figure S2B and Figure S2C). Importantly, models without admixture account for incomplete lineage sorting as they consider that populations share a common ancestry (different split times), and that each population has a specific effective size. To ensure models have the same number of parameters, in models without admixture we allowed for the possibility of a bottleneck associated with the split of *S. pyrenaicus* North, mimicking a founder effect. Models B and C without the possibility of bottlenecks were also considered. Second, we compared three other models to distinguish between a hybrid origin of *S. pyrenaicus* North and two models of secondary contact (Fig. S2D-F).

To obtain the observed minor allele frequency spectrum without missing data, we built the pairwise 2D-SFS by sampling 3 individuals from *S. carolitertii* and *S. pyrenaicus* South, and 4 individuals from *S. pyrenaicus* North sampled from Ocreza and Canha. We excluded Lizandro since it is an independent river basin, and its removal maximized the number of SNPs in the SFS. To sample individuals without missing data, we used the initial dataset without the MAF filter and divided it into blocks of 200bp. For each block, we sampled the individuals from each population with the least missing data keeping only the sites with data across all individuals. Since the SFS is affected by the depth of coverage, only genotypes with a depth of coverage >10x were used (Nielsen *et al*., 2011). This resulted in an observed SFS with 8 758 SNPs. For each model we performed 100 independent runs with 100 cycles, approximating the SFS with 100 000 coalescent simulations. Given that we used the SFS without monomorphic sites, all parameters were scaled in relation to a reference effective size, which was arbitrarily set to be the effective size *(Ne)* of *S. carolitertii*. To convert the relative divergence times estimated into absolute time in million years (Mya), we assumed a generation time of 3 years for these species (Magalhães *et al*., 2003; Almada and Sousa-Santos, 2010).

We then used the AIC to compare models. Since composite likelihoods computed with linked sites and pairwise 2D-SFS tend to overestimate the likelihood due to non-independence of data points (Excoffier *et al*., 2013), we built a 3D-SFS minimizing linked sites. We obtained 1 000 bootstrap joint 3D-SFS MAF by resampling one site from each GBS locus, assuming they are independent. We obtained the expected 3D-SFS for each model based on 100 000 coalescent simulations according to the parameters that maximized the composite likelihood. To account for noise of the approximation, we averaged the expected 3D-SFS of 10 runs. We then computed the likelihood of the 1 000 bootstrap unlinked 3D-SFS for each model. For each bootstrap replicate, we computed the AIC and relative likelihood of each model, following Excoffier *et al*., 2013.

### Detection of outlier loci in *S. pyrenaicus* North

To identify variation from the parental species (*S. carolitertii* or *S. pyrenaicus* South) that was likely selected in the introgressed *S. pyrenaicus* North we scanned for outlier loci using pairwise F_ST_ (Hudson *et al*., 1992). We assumed that selection favouring alleles from a target parental (P1) in the potential hybrid population (H) would lead to SNPs where (i) the target parental and hybrid have low differentiation (F_ST(H,P1)_ < quantile 0.05 F_ST(H,P1)_); (ii) the hybrid and the other parental (P2) have high differentiation (F_ST(H,P2)_ > quantile 0.95 F_ST(H,P2)_); and (iii) the differentiation between the hybrid and the other parental is higher than between the two parental populations (F_ST(H,P2)_ > quantile 0.95 F_ST(P1,P2)_ between parentals). This is because under neutrality we would expect the same level of differentiation between the minor parental (i.e., *S. pyrenaicus* South) and both the major parental (i.e., *S. carolitertii*) and the hybrid lineage (*S. pyrenaicus* North). Focusing on *S. carolitertii* and *S. pyrenaicus* (n=34), we applied a MAF filter of MAF>0.05 and kept only sites with more than 20% of data per population. We used only catalog loci with more than 2 SNPs and a mean distance between SNPs higher than 9. This analysis was repeated considering individuals from the three sampling locations of *S. pyrenaicus* North separately to identify outlier regions shared between them. Calculations were done in in RStudio v1.1.383 and R v3.4.4 using custom scripts. Finally, we blasted the identified outlier catalog loci against the NCBI database using the default settings of BLASTN (Zhang *et al*., 2000) and BLASTX (Altschul, 1997) v2.11.0+.

## Results

### Multispecies SNP dataset

After the initial processing, we obtained a mean of 5 759 264 high-quality reads per individual (Table S2). The catalog comprised 524 911 loci with a mean length of 160bp. Mean depth of coverage per sample was 59.7x. The filters applied resulted in 17 individuals with more than 50% of missing data, which were removed. The final dataset comprised 23 562 SNPs with 36.98% missing data and 48 individuals, as follows: *S. carolitertii* (n=10), *S. pyrenaicus* Ocreza (n=6), *S. pyrenaicus* Lizandro (n=4), *S. pyrenaicus* Canha (n=7), *S. pyrenaicus* Almargem (n=6), *S. pyrenaicus* Guadiana (n=1), *S. torgalensis* (n=9), and *S. aradensis* (n=5).

### Global patterns of genetic differentiation

Overall, the highest levels of genetic differentiation are between the two southwestern species (*S. torgalensis* and *S. aradensis*) and the two more widely distributed species (*S. carolitertii* and *S. pyrenaicus*) (F_ST_≤0.294; Table 1). The lowest levels of genetic differentiation were found between *S. pyrenaicus* North sampling locations and between them and *S. carolitertii* (F_ST_≤0.139). The latter were lower than between *S. pyrenaicus* North and *S. pyrenaicus* South (F_ST_≥0.171). Interestingly, the levels of differentiation found between both *S. carolitertii* and *S. pyrenaicus* North and *S. pyrenaicus* South are comparable to those found between *S. torgalensis* and *S. aradensis*.

**Table 1.** Pairwise F_ST_ calculated between the different sampling locations. *S. pyrenaicus* Guadiana was deliberately left out as there is only one individual from this sampling location.

|  | *S. carolitertii* | *S. pyrenaicus*<br>Ocreza | *S. pyrenaicus*<br>Lizandro | *S. pyrenaicus*<br>Canha | *S. pyrenaicus*<br>Almargem | *S. torgalensis* | *S. aradensis* |
| --- | --- | --- | --- | --- | --- | --- | --- |
| *S. carolitertii* | - | 0.101 | 0.139 | 0.060 | 0.186 | 0.320 | 0.310 |
| *S. pyrenaicus* Ocreza | - | - | 0.138 | 0.054 | 0.204 | 0.344 | 0.331 |
| *S. pyrenaicus* Lizandro | - | - | - | 0.075 | 0.238 | 0.372 | 0.357 |
| *S. pyrenaicus* Canha | - | - | - | - | 0.171 | 0.309 | 0.294 |
| *S. pyrenaicus* Almargem | - | - | - | - | - | 0.351 | 0.338 |
| *S. torgalensis* | - | - | - | - | - | - | 0.184 |
| *S. aradensis* | - | - | - | - | - | - | - |

The first three principal components (PCs) of the PCA explain ~25% of the variation (Figure S3). PC1 (Figures 2A.1 and S3C) explains ~14% of the variance and clearly separates two groups: (i) *S. carolitertii* and *S. pyrenaicus*, and (ii) *S. aradensis* and *S. torgalensis*. This is consistent with the higher pairwise F_ST_ values obtained between these two groups. PC2 explains ~6% of the variance and separates *S. aradensis* from *S. torgalensis* (Figure 2A). Finally, PC3 explains ~5% separates *S. pyrenaicus* South from a cluster formed by *S. carolitertii* and *S. pyrenaicus* North (Figure 2A.2 and S3C).

**Figure 2.**
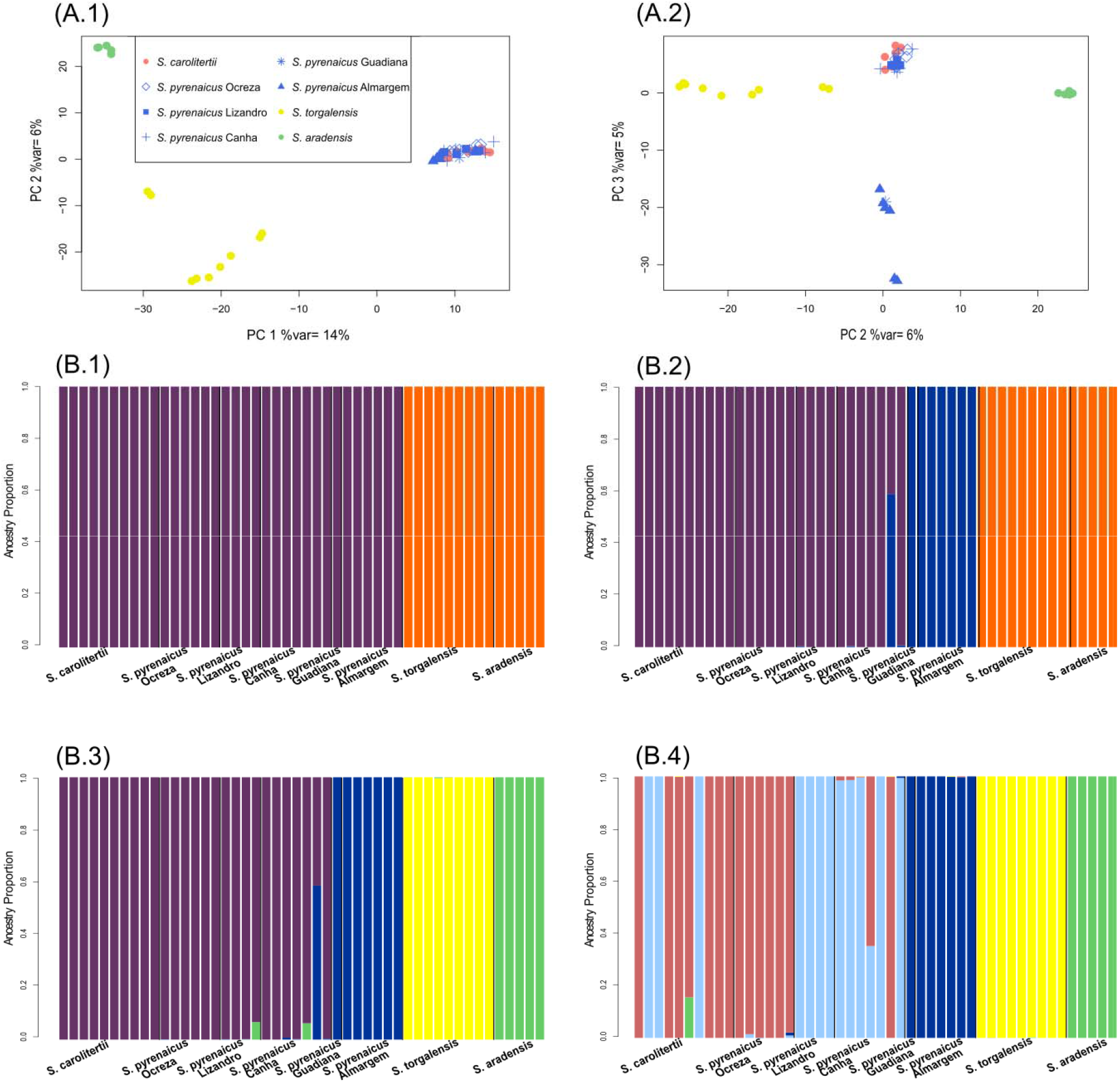
Global patterns of genetic differentiation. **(A) Principal Component Analysis.** (A.1) PC1 and PC2; (A.2) PC2 and PC3. Each point corresponds to one individual. **(B) Individual ancestry proportions inferred with ADMIXTURE.** (B.1) K=2; (B.2) K=3; (B.3) K=4; (B.4) K=5. Each vertical bar corresponds to one individual and the proportion of each colour corresponds to the estimated ancestry proportion from a given cluster. Individuals are grouped from north to south. Sampling locations separated by black lines.

Regarding the ADMIXTURE analysis, K=2 achieved the lowest cross-validation (Figure S4). The results were consistent across the 100 runs for 2≤K≤5 (mean G’=1.0 for K=2, K=4 and K=5 and mean G’≥0.9974 for K=3). For the best value of K (K=2), one of the clusters includes the two southwestern species (*S. torgalensis* and *S. aradensis)*, while the other comprises *S. carolitertii* and *S. pyrenaicus* (Figure 2B.1). This clustering mimics the separation created by PC1 in the PCA. For K=3, a third cluster formed by *S. pyrenaicus* South appears (Figure 2B.2). For K=4 the two southwestern species (*S. torgalensis* and *S. aradensis)* are differentiated into different clusters (Fig 2B.3). Interestingly, in these last two K values, one of the individuals from the Canha sampling location of *S. pyrenaicus* North shows a significant proportion from the *S. pyrenaicus* South cluster. Two *S. pyrenaicus* North individuals also show a small proportion of the *S. aradensis* cluster in K=4, which may be due to ancestral polymorphism. K=5 does not fit the data as well (mean cross validation much higher than for 2≤K≤4, Figure S4), and fails to separate *S. carolitertii* from *S. pyrenaicus* North.

### Relationship between species and populations

The unrooted population tree obtained with TreeMix (Figure 3) shows a clear separation between two groups: (i) *S. aradensis* and *S. torgalensis*, and (ii) *S. carolitertii* and *S. pyrenaicus*. Within the group of *S. carolitertii* and *S. pyrenaicus*, we found two main lineages: *S. pyrenaicus* South (here represented by *S. pyrenaicus* Almargem) and *S. carolitertii* and *S. pyrenaicus* North, which is consistent with pairwise F_ST_, PCA and ADMIXTURE results. One of the two migration edges inferred is between *S. pyrenaicus* North and *S. pyrenaicus* South.

**Figure 3.**
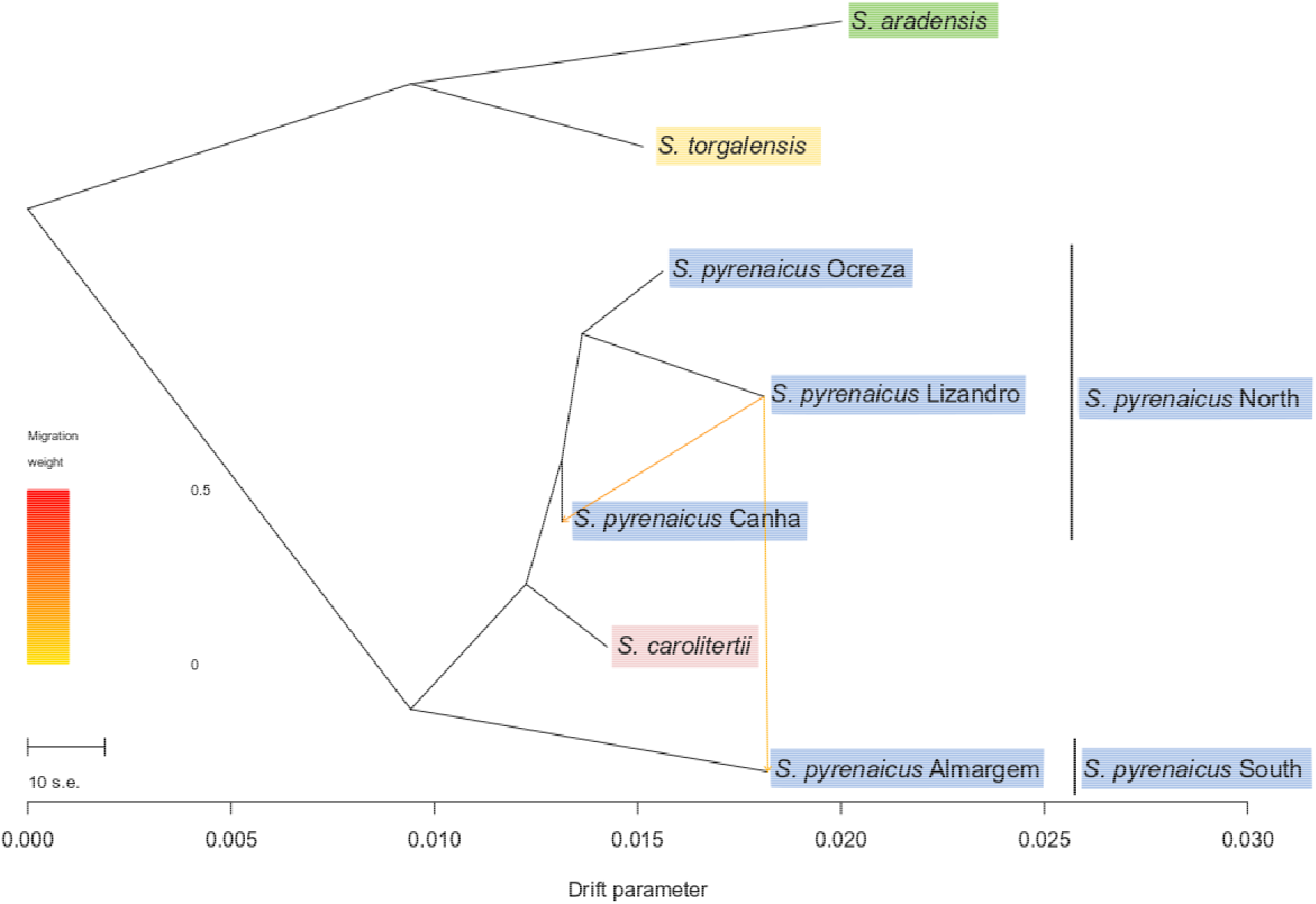
Unrooted species tree graph obtained with TreeMix. Branch lengths are represented in units of genetic drift, i.e., the longer a given branch the stronger the genetic drift experienced in that lineage. Higher genetic drift can be due to older divergence times or smaller effective population sizes of a given lineage. Arrows represent migration events.

Regarding the maximum likelihood phylogeny of individuals (Figure S5), the topology of the tree is consistent with the one obtained for the populations with TreeMix, showing two well supported clades. It strongly supports the paraphyly of *S. pyrenaicus* with respect to *S. carolitertii*, with two well supported subclades with individuals of: (i) *S. carolitertii* and *S. pyrenaicus* North, and (ii) *S. pyrenaicus* South.

### Effect of linked SNPs

The dataset with one SNP per block of 200bp comprised 2 607 SNPs with ~32.4% missing data. The results of PCA, ADMIXTURE and TreeMix analysis were overall consistent with those from the initial dataset of 23 562 SNPs (Figures S6-S8).

### Detection of introgression between *S. carolitertii* and *S. pyrenaicus*

When considering *S. carolitertii* as the potential source of introgression, we found significant positive D-statistic values indicating an excess of sites where *S. pyrenaicus* North shares the same allele with *S. carolitertii*, independently of the outgroup used (Figure 4A). This could indicate that *S. carolitertii* and *S. pyrenaicus* North share a more recent common ancestor, in agreement with TreeMix and phylogeny results (Figures 3 and S5). However, when considering *S. pyrenaicus* South as the potential source of introgression we also found significant positive D-statistic values, indicating an excess of sites where both *S. pyrenaicus* share the same allele (Figure 4B), in agreement with the TreexMix migration edge (Figure 3). This indicates that the relationship between the species is not described by a bifurcating tree with a more recent common ancestry of *S. carolitertii* and *S. pyrenaicus* North, as we would expect D=0 if that was the case.

**Figure 4.**
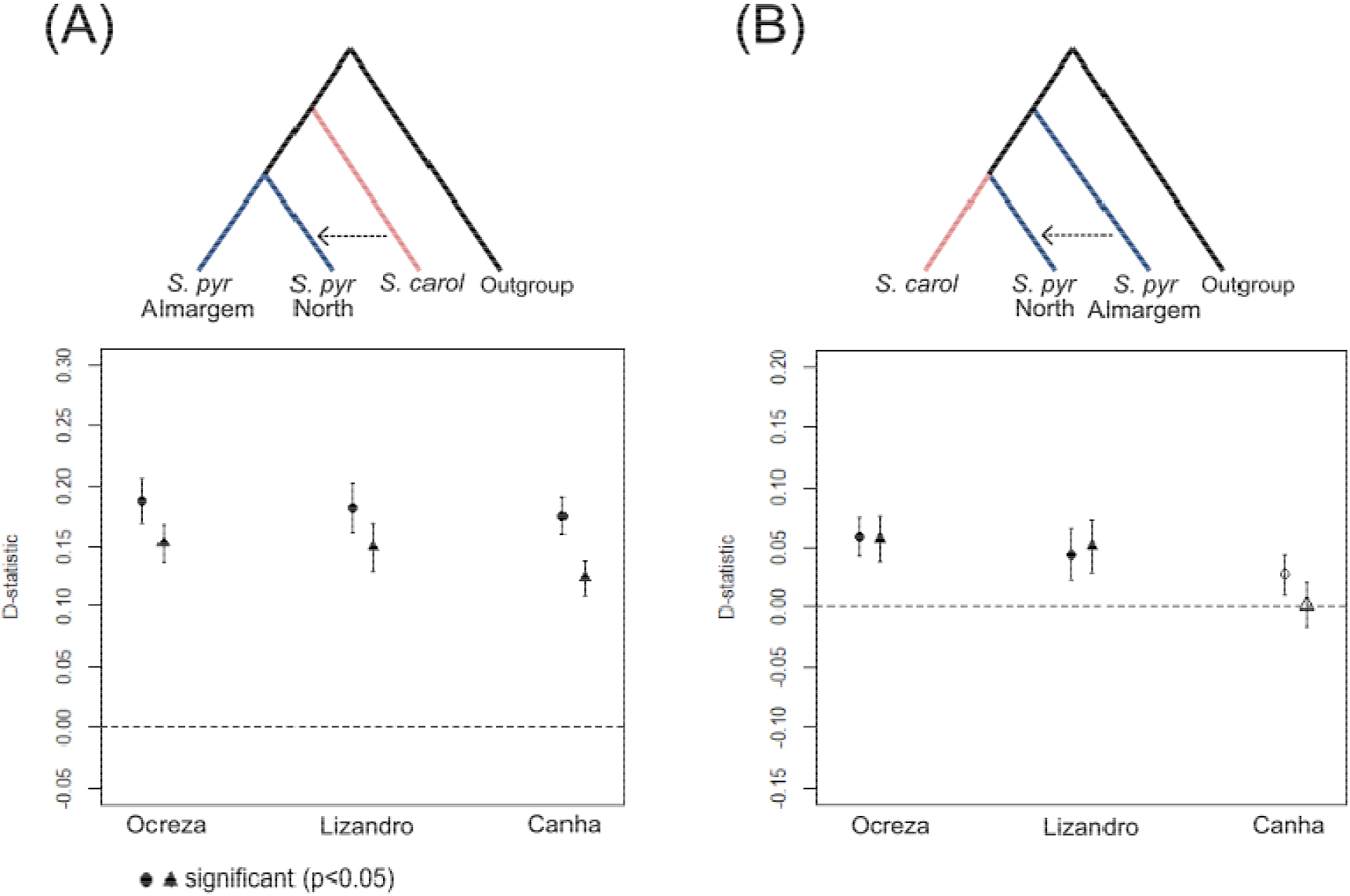
D-statistics calculated for different topologies. For each topology, the results are shown for the different *S. pyrenaicus* North sampling locations (Ocreza, Lizandro, Canha) used. “S. carol” stands for *S. carolitertii* and “S.pyr Almargem” stands for *S. pyrenaicus* Almargem. Results obtained with each outgroup are represented by a different symbol (circles for *S. torgalensis* and triangles for *S. aradensis)*. Full symbols represent significant D values (p<0.05).

We find no evidence of a geographical cline of admixture with *S. carolitertii* along the *S. pyrenaicus* North distribution (Figure S9), with its northern most sampling location (Ocreza – location 2 on Figure 1) showing no signs of sharing more alleles with *S. carolitertii* than with other *S. pyrenaicus* North populations. Thus, the inexistence of a geographical cline of admixture and the consistent signal across different *S. pyrenaicus* North sampling locations in Figure 4 suggest a scenario of introgression between *S. carolitertii* and the ancestor of *S. pyrenaicus* North prior to the divergence of the different *S. pyrenaicus* North populations.

In the case of recent introgression events, we would expect to find differences in the D-statistic among individuals from a given population. However, when we computed the D-statistic by individual (Figure S10 and Table S4), we found limited variation among different individuals from the same population, suggesting that introgression events are likely pre-dating the divergence of populations.

### Demographic modelling of divergence of *S. carolitertii* and *S. pyrenaicus*

First, we compared models with admixture (Figure 5A) to models of bifurcating trees without gene flow accounting for incomplete lineage sorting, i.e. different topologies, times of split and effective population sizes (Figures 5B-C). The model with admixture reached a higher likelihood (Table S5), approximately 9.00 log10 units higher than the best bifurcating tree model with the topology supported by the individual-based phylogeny (Figure S5), i.e., with a more recent ancestor of *S. carolitertii* and *S. pyrenaicus* North (Figure 5B). This result was not affected by the number of parameters, as bifurcating tree models with or without an extra effective size parameter to allow for a potential bottleneck (see Methods) had lower likelihood values than the admixture model (Tables S5, S6A, S7 and S8). This supports that *S. pyrenaicus* North likely experienced introgression and that its relationship with *S. carolitertii* and *S. pyrenaicus* South cannot be explained by a simple bifurcating tree, in agreement with TreeMix and D-statistics (Figures 3–4). Next, we explored models to distinguish between a scenario of hybrid origin of *S. pyrenaicus* North (Figure 5D) and secondary contact (Figure 5D-F). The parameter estimates were similar and consistent across models with admixture (Figure 5A, D-F), indicating (i) a major contribution of 80-84% from *S. carolitertii* into *S. pyrenaicus* North; (ii) a relatively old divergence of the parental lineages (~15 times older than admixture times); and (iii) relatively larger effective sizes in ancestral lineages (Figures 5A, 5D-F). To convert relative parameters into years we considered different effective sizes of *S. carolitertii*, which was considered as a reference (Tables S6A-B). To compare models, we computed the relative likelihoods based on AIC for 1000 bootstrap datasets with a single SNP per GBS locus. Model D achieved the higher relative likelihood, irrespective of the threshold used to pool SFS entries (Figure S11). The best model D is compatible with a “hybrid origin”, inferring that at time of admixture *S. pyrenaicus* North received ~84% from *S. carolitertii* (major parental) and the remaining ~16% from *S. pyrenaicus* South (minor parental).

**Figure 5.**
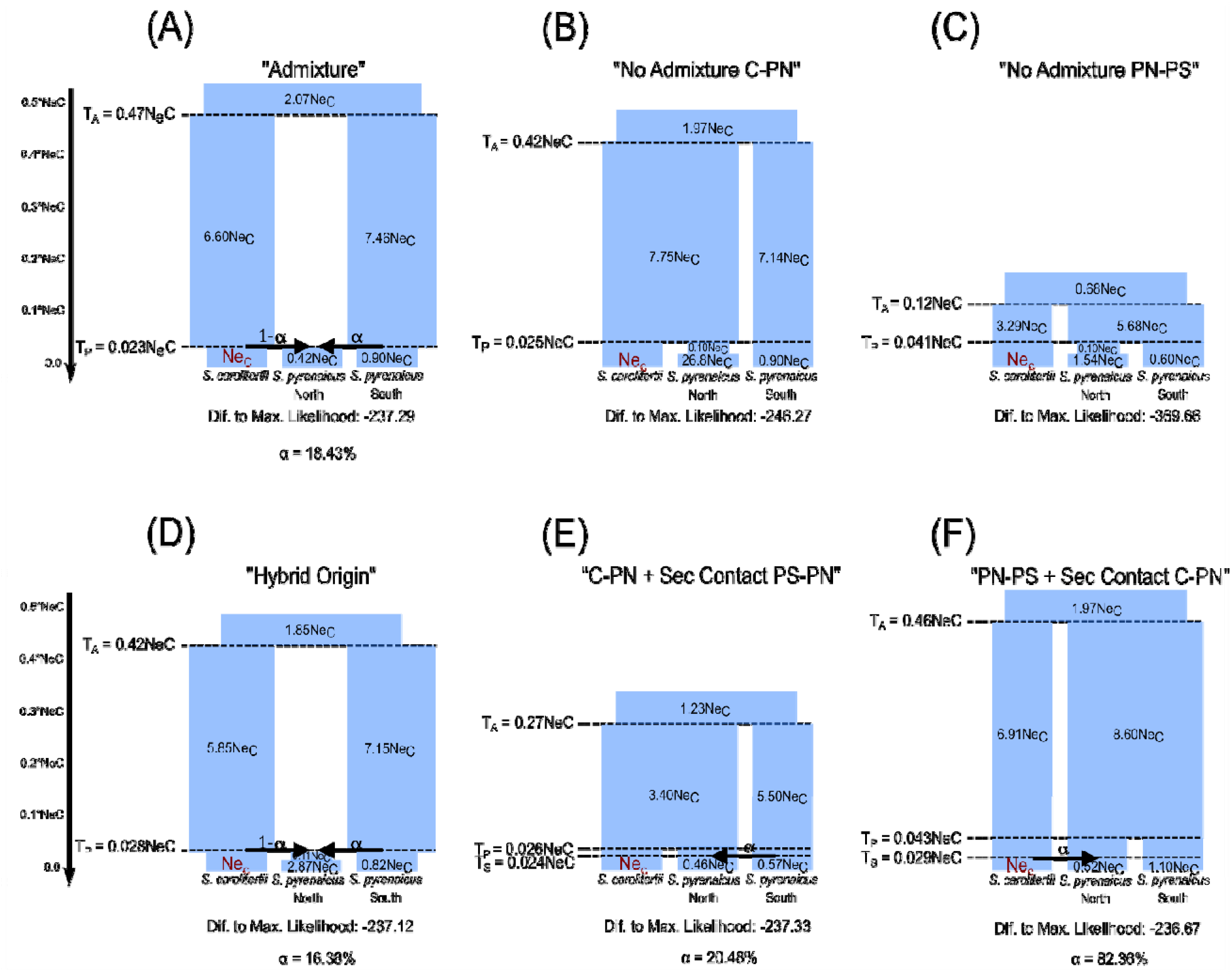
Schematic representation of the likelihood of the models tested with *fastsimcoal2*. The name given to each model is indicated above the schematic representation, as well the difference to maximum likelihood (Dif. To Max. Likelihood) which is the difference in log10 units between the estimated likelihood and the maximum likelihood if there was a perfect fit to the observed site frequency spectrum. The closer to zero (less negative values), the better the fit. The estimated admixture proportion is the parameter α. Models (A) to (C) have 8 parameters and therefore are directly comparable. Models (D) to (F) have 9 parameters and are also directly comparable. All inferred parameters are indicated in relation to the Ne of *S. carolitertii* (NeC)). T_A_ – divergence from the ancestral; T_P_ – divergence of the northern *S. pyrenaicus*; T_S_ – secondary contact. Population sizes are not to scale. See supplementary Figure S11 for relative likelihoods based on AIC, and Supplementary Tables S5-S8 for estimated likelihoods and parameter values.

### Detection of outlier loci in *S. pyrenaicus* North

We identified 10 outlier loci with low F_ST_ between *S. carolitertii* and *S. pyrenaicus* North and high F_ST_ between *S. pyrenaicus* North and *S. pyrenaicus* South (Figure 6A – red triangles). These correspond to alleles from *S. carolitertii* which might have been selected in *S. pyrenaicus* North. However, when separating *S. pyrenaicus* North into the three sampling locations, none of these was shared between them (Figure 6B). Moreover, none produced relevant BLAST hits.

**Figure 6.**
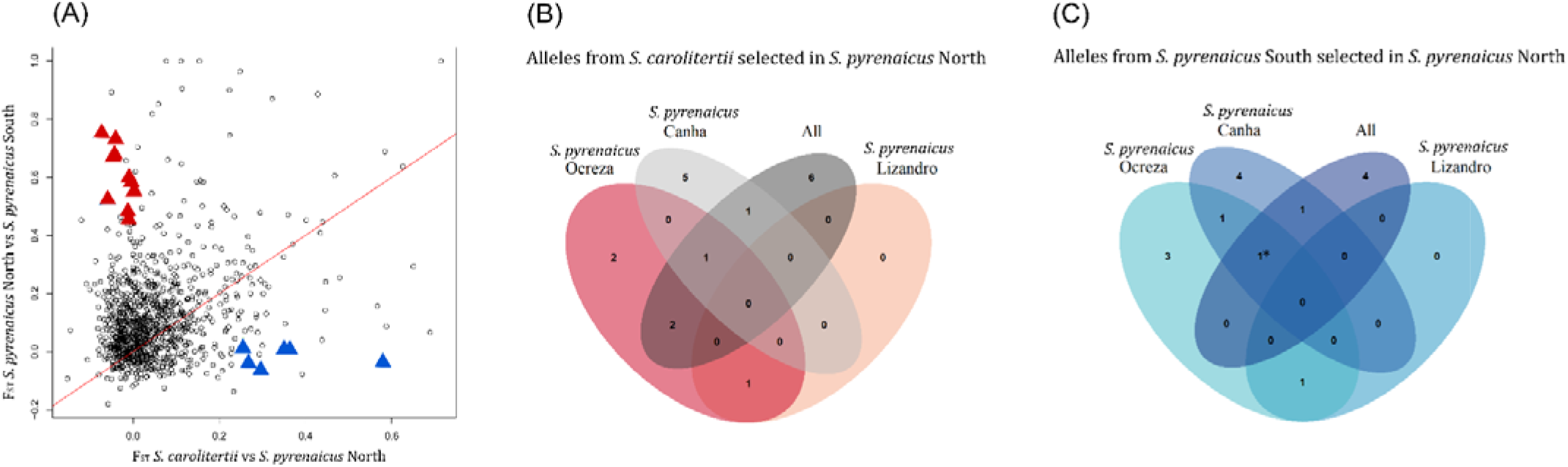
**(A) Outlier loci detected in *S. pyrenaicus* North.** Red triangles correspond to alleles from *S. carolitertii* potentially selected for in *S. pyrenaicus* North and blue triangles to alleles from *S. pyrenaicus* South potentially selected in *S. pyrenaicus* North. **(B). Outlier loci from *S. carolitertii* shared between the different *S. pyrenaicus* North sampling locations (C). Outlier loci from *S. pyrenaicus* South shared between the different *S. pyrenaicus* North sampling locations.** Asterisk indicates the catalog loci identified as *vitellogenin*.

We also identified 6 loci with high F_ST_ between *S. carolitertii* and *S. pyrenaicus* North and low F_ST_ between *S. pyrenaicus* North and *S. pyrenaicus* South (Figure 6A – blue triangles). These correspond to alleles from *S. pyrenaicus* South that might have been selected in *S. pyrenaicus* North. One of the identified catalog loci, shared between the individuals from the locations “Ocreza” and “Canha” and also identified when all locations were pooled together (Fig 6C), revelated an alignment match to *vitellogenin*, mostly from other cyprinid fish (Table S9). The other 5 outlier loci produced no relevant BLAST hits.

## Discussion

### Contrasting role of gene flow in the two species’ pairs

The genomic analysis presented here indicates a species tree comprising two main groups: (i) the one of *S. torgalensis* and *S. aradensis* and (ii) that of *S. carolitertii* and *S. pyrenaicus*. This is in agreement with phylogenies previously obtained for mitochondrial (Brito *et al*., 1997; Sanjur *et al*., 2003; Mesquita *et al*., 2007) and different nuclear markers (Almada and Sousa-Santos, 2010; Waap *et al*., 2011; Sousa-Santos *et al*., 2019; Perea *et al*., 2020). The results also show strong genetic differentiation between the two species’ pairs (Table 1, Figures 2–3), as formerly reported (Coelho *et al*., 1995; Almada and Sousa-Santos, 2010), which likely reflects their old divergence time, estimated to be approximately 14Mya based on phylogenies of mitochondrial and nuclear genes (Perea *et al*., 2010; Sousa-Santos *et al*., 2019).

Gene flow appears to have played a very different role on the evolutionary history of the two specie’s pairs. We found no evidence of gene flow involving *S. torgalensis* and *S. aradensis*. While we could not calculate D-statistics due to our sampling scheme, the results of the analyses including the four species (Table 1, Figures 2–3) did not support interspecific gene flow involving *S. torgalensis* and/or *S. aradensis*. Thus, for these two species with more restricted distribution areas our results are compatible with divergence of isolated lineages in allopatry. In contrast, for the two species with wider distributions (*S. carolitertii* and *S. pyrenaicus)* our results indicate gene flow was paramount in their evolutionary history, specifically in the *S. pyrenaicus* North lineage (Figures 4–5). This contrasting role of gene flow between the two more restricted and the two more widely distributed species might reflect their historical dispersal routes into different river basins. Our results are consistent with the hypothesis that the ancestor of the two southwestern species (*S. torgalensis* and *S. aradensis)* became isolated in southwestern Iberia around 11.6-7.2Ma, while *S. carolitertii* and *S. pyrenaicus* only reached their current ranges in the last 2.6Ma (Sousa-Santos *et al*., 2019), the latter as a consequence of geological changes that led to the current conformation of the major Iberian river basins (Cunha *et al*., 2005; Casas-Sainz and de Vicente, 2009; Pais *et al*., 2012; Antón *et al*., 2014). Thus, the older isolation of river basins in southwestern Iberia probably created less opportunity for gene flow involving *S. torgalensis* and *S. aradensis*. This is in agreement with results from other freshwater fish species (Sousa *et al*., 2010) and amphibians (Martínez-Solano *et al*., 2006; Gonçalves *et al*., 2009), indicating higher genetic differentiation in this geographic region.

### Evidence of hybridization between *S. carolitertii* and *S. pyrenaicus*

Our demographic modelling results indicates that the *S. pyrenaicus* North lineage is likely the result of hybridization – the best model estimates indicate 84% contribution from *S. carolitertii* and the remaining 16% from *S. pyrenaicus* South (Figure 5D and Figure S12). A hybridization scenario is also supported by other analyses (Figures 3–4). For example, TreeMix infers a common ancestor for *S. carolitertii* and *S. pyrenaicus* North, together with a migration event between northern and southern *S. pyrenaicus*, indicating allele sharing that cannot be explained by a bifurcating tree (Figure 3). The uncovered hybridization provides an explanation for the incongruences between mitochondrial and nuclear markers reported in previous studies: while in mtDNA phylogenies all *S. pyrenaicus* had a more recent common ancestor (Brito *et al*., 1997; Mesquita *et al*., 2007), in nuclear phylogenies individuals of river basins corresponding to *S. pyrenaicus* North shared a more recent ancestor with *S. carolitertii* (Waap *et al*., 2011; Sousa-Santos *et al*., 2019; Perea *et al*., 2020).

The D-statistics results indicate that the hybridization event predates the differentiation of the three *S. pyrenaicus* North sampling locations, since the introgression signal is shared between them (Figure 4). Given that Lizandro (Figure 1 – location 3) is an independent river basin with no connection to the Tagus basin (to which the other two *S. pyrenaicus* North sampling locations belong), the most likely explanation is that the hybridization event predates the colonization of this independent river. Thus, the combination of the D-statistics with the best demographic model indicates that *S. pyrenaicus* North likely originated through hybridization and posteriorly dispersed to its current range. Geological data indicates that prior to reaching their current conformation, the major Iberian river basins were endorheic, with the transition to exorheism occurring in the last 2.6Ma (Cunha *et al*., 2005; Casas-Sainz and de Vicente, 2009; Pais *et al*., 2012; Antón *et al*., 2014). Thus, it is possible that this transition increased the chances of contact between previously isolated lineages. Another possibility is that these “lake like” endorheic basins allowed for lineages to mix, with posterior dispersal of resulting hybrids.

The origin of *S. pyrenaicus* North through a single hybridization event followed by isolation, as modelled in Figure 5D, is consistent with a scenario of hybrid speciation. Most authors agree that homoploid hybrid speciation (HHS), or recombinational speciation, occurs when hybridization plays a key role in establishing a new reproductively isolated lineage without changes in ploidy (Anderson and Stebbins, 1954; Grant, 1981; Mallet, 2007; Mavárez and Linares, 2008; Abbott *et al*., 2013). Schumer *et al*., 2014 establish three criteria to demonstrate HHS: (1) hybrid lineages must be reproductively isolated from the parental ones that originated them; (2) the genome of the hybrids must show evidence of hybridization; (3) reproductive isolation must be demonstrated to be a consequence of hybridization. Our results demonstrate *S. pyrenaicus* North meets the second criterion. Additionally, it has a non-overlapping distribution with *S. carolitertii* and *S. pyrenaicus* South, thus being currently reproductively isolated from its parentals through their allopatric distributions, and ploidy is maintained between the three lineages (2n=50) (Collares-Pereira *et al*., 1998). Yet, we currently lack information on the possible role of hybridization on the establishment of reproductive isolation to meet the three criteria. Indeed, very few biological systems clearly meet all the criteria outlined above. A notable example are the *Helianthus* sunflowers, in which hybridization between two species produced independent linages reproductively isolated from the parentals, with clear evidence of hybridization in their genomes, that can occupy dry and saline environments that the parents cannot, as a direct result of hybridization (Rieseberg *et al*., 1995, 1996, 2003). In freshwater fish, some taxa have been proposed as being the result of HHS (e.g. DeMarais *et al*., 1992), although prior to the definition of the above criteria.

### Potential implications of hybridization in *S. pyrenaicus* North

We estimated relative times indicating that the parental *S. carolitertii* and *S. pyrenaicus* South lineages diverged much earlier than the hybridization event that originated the *S. pyrenaicus* North lineage, inferring that the time of hybridization is 0.07 of the divergence time (Figure 5D, Table S6B). In this scenario, *S. pyrenaicus* North might have benefited from the combination of different alleles from the two parentals, already filtered out by selection over many generations. Yet, the accumulation of genetic differentiation between the two parental lineages may also have led to genetic incompatibilities when the two genomes mixed. Our estimates indicate an asymmetric and larger contribution from *S. carolitertii* at the time of hybridization (~84%). Such asymmetries in admixture proportions have been reported in other freshwater fish, albeit in cases of recent hybridization, e.g., swordtail fish (Schumer *et al*., 2018) and sticklebacks (Marques, Lucek, *et al*., 2019). Here, this asymmetry could reflect the relative proportions of *S. carolitertii* and *S. pyrenaicus* South in the initial reproductive pool that originated *S. pyrenaicus* North. Another possibility is that incompatibilities might have been resolved towards the *S. carolitertii* lineage. Notably, at mitochondrial markers *S. pyrenaicus* North is more closely related with *S. pyrenaicus* South (Brito *et al*., 1997; Mesquita *et al*., 2007), despite the larger nuclear contribution of *S. carolitertii*. This can be due to neutral stochasticity, but it might also be related to cito-nuclear incompatibilities or behavioural mechanisms favouring mating between *S. carolitertii* males and *S. pyrenaicus* South females.

Despite the asymmetry in the contribution of both parental lineages, there is variation in differentiation along the genome between the hybrid and parental lineages (Figures 6 and S12). This variation reflects the stochasticity of neutral processes (e.g., drift and gene flow), but it also provides indirect evidence for the action of selection removing incompatibilities and/or favouring specific parental alleles (adaptive introgression). We identified 6 outlier regions compatible with selection of alleles from the minor parental lineage in *S. pyrenaicus* North, one of them corresponding to *vitellogenin*. Vitellogenin is an egg yolk precursor that is usually only expressed in females during oogenesis (Tyler *et al*., 1996), although its expression can also be triggered in males of different fish species in response to exogenous oestrogen exposure (Harries *et al*., 1997; Flammarion *et al*., 2000; Van Den Belt *et al*., 2003). Given its role in egg yolk formation, *vitellogenin* is of great importance to egg-laying organisms like fish. One possibility is that this genomic region was selected and maintained from the minor parental (*S. pyrenaicus* South) due to adaptive introgression, although the underlying selective pressures remain unclear. However, we cannot discard the hypothesis that genetic incompatibilities were involved and thus constrained the maintenance of this region from *S. pyrenaicus* South.

While we could not annotate other outlier genomic regions of interest in *S. pyrenaicus* North, the geographic distribution of this lineage at an intermediate between the Atlantic and Mediterranean climate types provides future opportunity to investigate the potential adaptive role of hybridization. In particular, increasing temperatures and propensity for seasonal drought from north to south (the latter leading to isolation of the fish in the deeper water sections during the summer, when large portions of the riverbed dry out) (Gasith and Resh, 1999; Magalhães *et al*., 2003; Jesus *et al*., 2017) might impose strong selective pressures. This is consistent with previous studies that found (i) signatures of positive selection based on dN/dS on circadian genes likely associated with thermal adaptation in *Squalius* species (Moreno *et al*., 2021), and (ii) differences in gene expression between species inhabiting Atlantic (*S. carolitertii)* and Mediterranean climates (*S. torgalensis)* when exposed to increasing temperatures (Jesus *et al*., 2016, 2017). Thus, its hybrid origin might confer some advantage to *S. pyrenaicus* North at intermediate environmental conditions, as described in other organisms. For example, in *Saccharomyces* yeast laboratory crosses between *S. cerevisiae* and *S. paradoxus* produced transgressive F2 hybrids that slightly increased their environmental range and outcompeted their parentals across the cline (Stelkens *et al*., 2014). In freshwater fish, hybrids between low and high elevation swordtail species experience higher fitness at intermediate conditions (Culumber *et al*., 2012).

### Implications for conservation

*S. pyrenaicus* populations, like other endemic Iberian cyprinids, have been impacted by habitat degradation, due to the construction of dams and water extraction for irrigation, and the introduction of exotic species. Consequently, *S. pyrenaicus* is listed as “Endangered” in the Portuguese Red List (Cabral *et al*., 2005) and as “Vulnerable” in the Spanish Red List (Doadrio, 2002). Our work provides evidence of two very distinct gene pools, each with its own evolutionary history, within the analysed *S. pyrenaicus* distribution: the one inhabiting the Tagus and small Atlantic coastal basins (*S. pyrenaicus* North) and the one distributed along the Guadiana and small Mediterranean basins (*S. pyrenaicus* South). Thus, this difference should be considered in future management efforts.

## Conclusions

Our study revealed a species tree with two well differentiated groups in which gene flow had a contrasting role: (i) *S. torgalensis* and *S. aradensis* and (ii) *S. carolitertii* and *S. pyrenaicus*. We find no evidence of past gene flow in the first, consistent with divergence in allopatry. In contrast, we uncover past hybridization between *S. carolitertii* and *S. pyrenaicus*, originating the *S. pyrenaicus* North lineage, with ~84% contribution from *S. carolitertii* and ~16% from *S. pyrenaicus* South. Our estimates indicate that a single hybridization event followed by isolation, consistent with a scenario of hybrid speciation, is more likely than a secondary contact scenario. However, the scenario might be more complex, e.g., involving further changes in the past effective sizes. In the future, whole genome sequencing data and more extensive sampling would be helpful to explore the sources of the unbalanced estimated admixture proportions in *S. pyrenaicus* North and evaluate the adaptive potential of this hybridization, given the contrasting environments of the parental lineages.

This work adds to the growing list of examples where hybridization has been uncovered, describing a study system suitable for future work on the processes and consequences of hybridization. Much remains to be understood regarding how hybridization shapes the genomes of species and genetic incompatibilities are resolved, its relation to the evolution of reproductive barriers between hybrid and parental lineages, and its role in adaptation.

## Acknowledgements

We thank Tiago F. Jesus for the help in the preparation of the samples. This work was funded by the strategic projects UID/BIA/00329/2013 (2015-2018) and UIDB/00329/2020 granted to cE3c from the Portuguese National Science Foundation - Fundação para a Ciência e a Tecnologia (FCT). SLM is funded by an FCT scholarship (SFRH/BD/145153/2019). VCS was funded by FCT (CEECIND/02391/2017 and CEECINST/00032/2018/CP1523/CT0008) and by EU H2020 program (Marie Skłodowska-Curie grant 799729). We thank the INCD (https://incd.pt/) for use of their computing infrastructure, which is funded by FCT and FEDER (project 01/SAICT/2016 n° 022153).

## Conflict of interest

The authors declare that they have no conflict of interest.

## Notes

### Competing Interest Statement

The authors have declared no competing interest.

### Summary of Updates

We have modified the bioinformatic pipeline to minimize missing data, and have repeated all downstream analyses. A new author was added due to his contribution in these analyses. The new version of the manuscript have the updated results with major changes in the text and figures. Yet, the results are qualitatively the same as in the previous version, but several results have been updated and new analyses have been included in the manuscript.

