## Supplementary Figures for "Genomic data and multi-species demographic modelling uncover past hybridization between currently allopatric freshwater species"

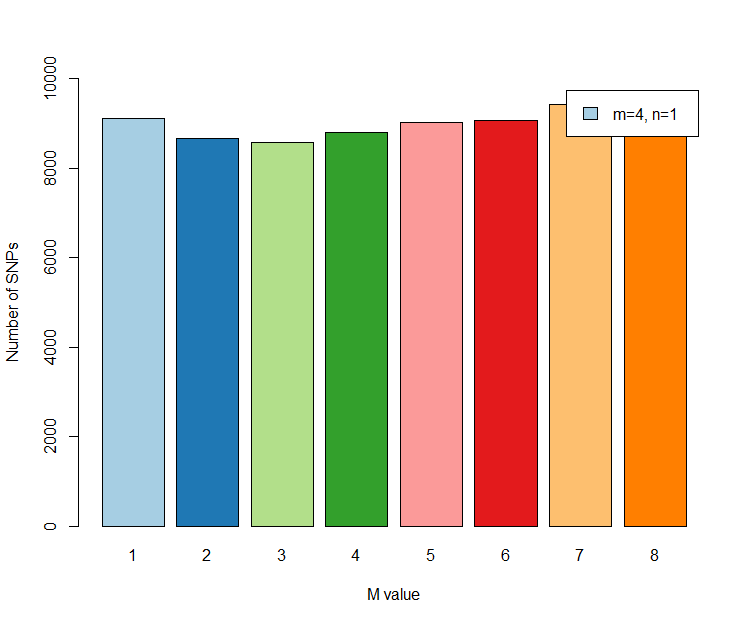

**(A)**

**Figure S1 - Number of SNPs obtained with different M and n values.** **(A)** The number of mismatches allowed between sequences from the same individual (M) was varied from 1 to 8 while all the other parameters were kept fixed (m=4, n=1). **(B)** The number of mismatched allowed between sequences from different individuals (n) was varied from 1 to 9 while all the other parameters were kept unchanged (m=4, M=2). In both cases, SNPs were required to be present in all populations in 50% of the individuals.

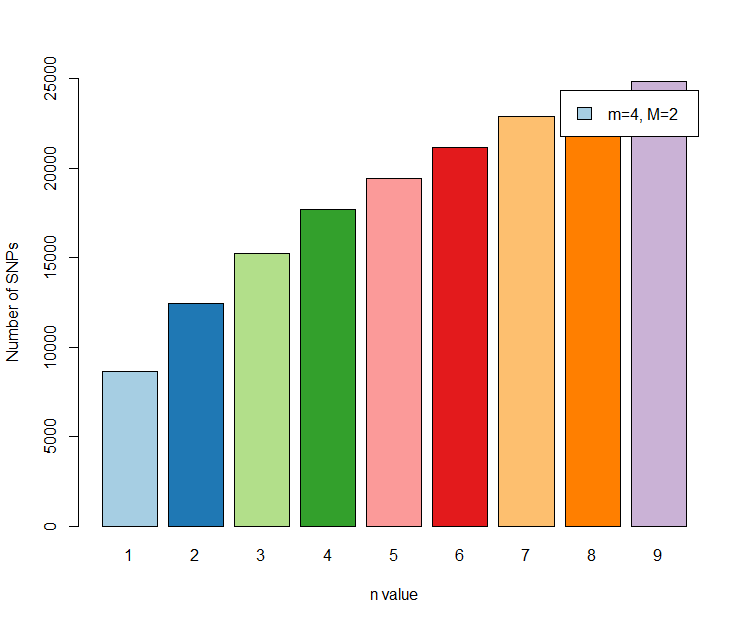

**(B)**

**Figure S1 (cont.) - Number of SNPs obtained with different M and n values.** **(A)** The number of mismatches allowed between sequences from the same individual (M) was varied from 1 to 8 while all the other parameters were kept fixed (m=4, n=1). **(B)** The number of mismatched allowed between sequences from different individuals (n) was varied from 1 to 9 while all the other parameters were kept unchanged (m=4, M=2). In both cases, SNPs were required to be present in all populations in 50% of the individuals.

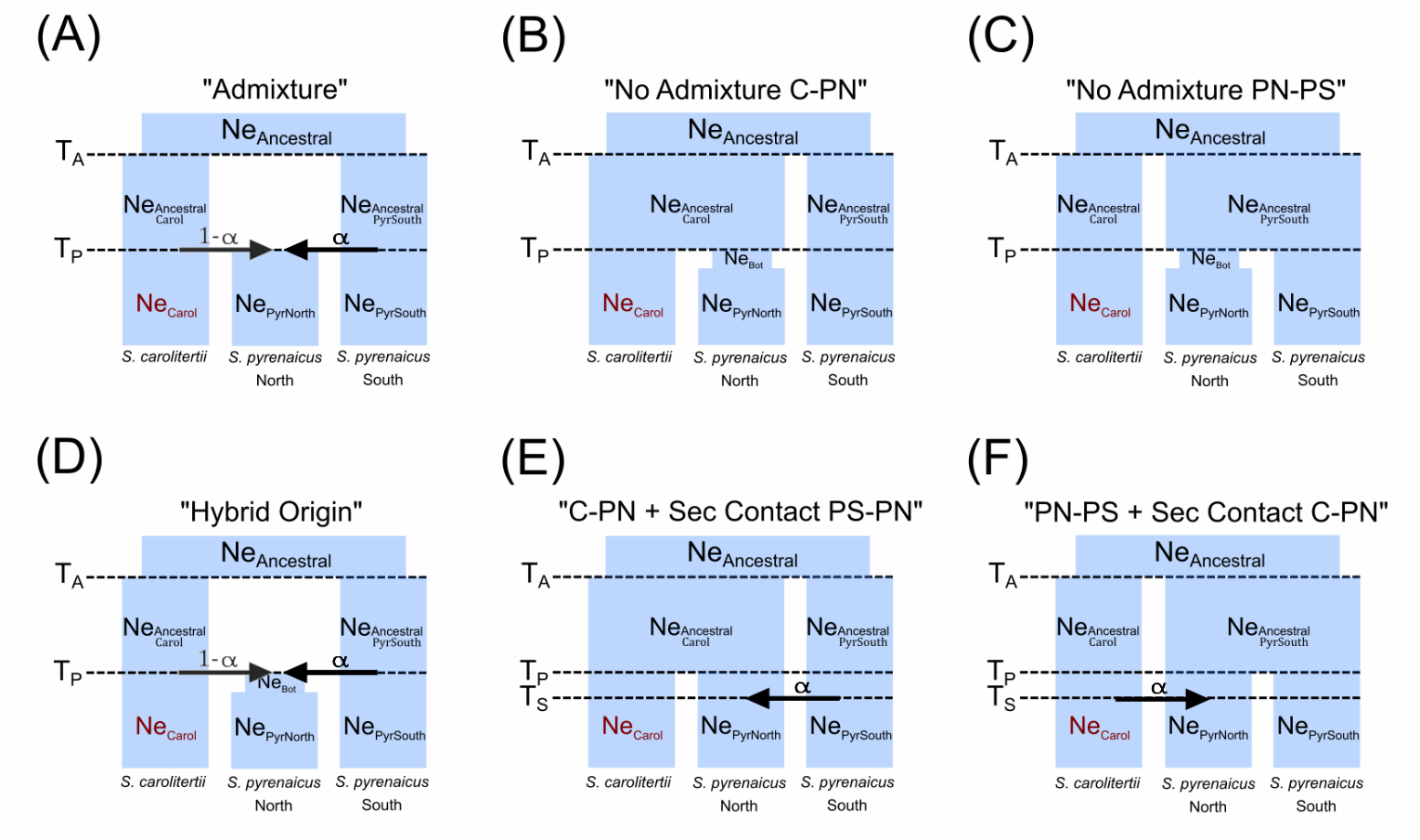

**Figure S2 – Models compared and parameters inferred for each model with fastsimcoal2.** Models (A) to (C) have the same number of parameters (number_parameters_ = 8) and the same is true for models (D) to (F) (number_parameters_ = 9). In all models Ne_Carol_ is coloured in red to indicate this is not an inferred parameter but rather the value in reference to which all other parameters are inferred. In models (A) and (D), the contribution of S. carolitertii into S. pyrenaicus North is not directly inferred (and thus is coloured in dark grey) but obtained simply by subtracting the estimated contribution of S. pyrenaicus South (1-α).

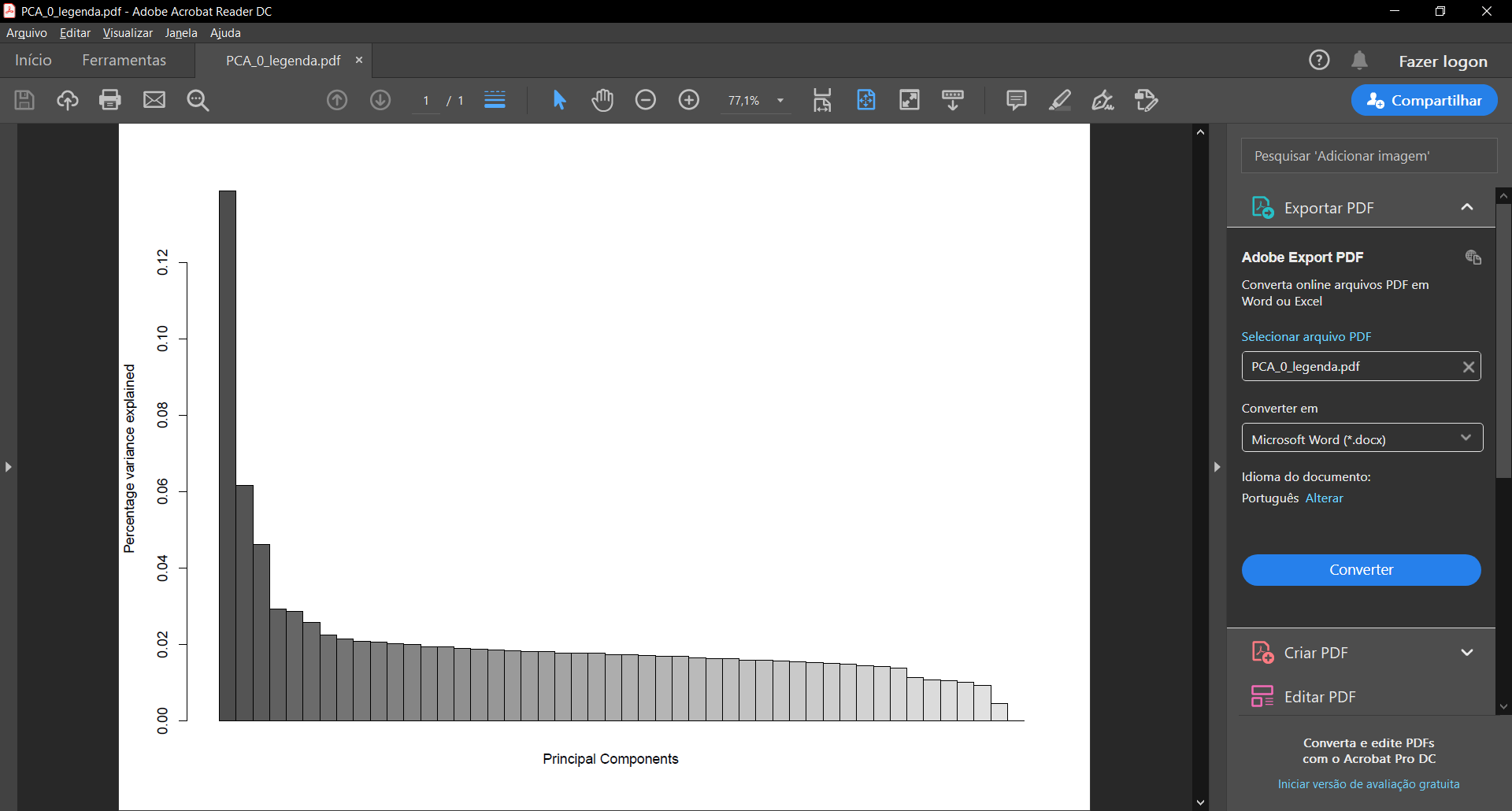

**(A)**

**(B)**

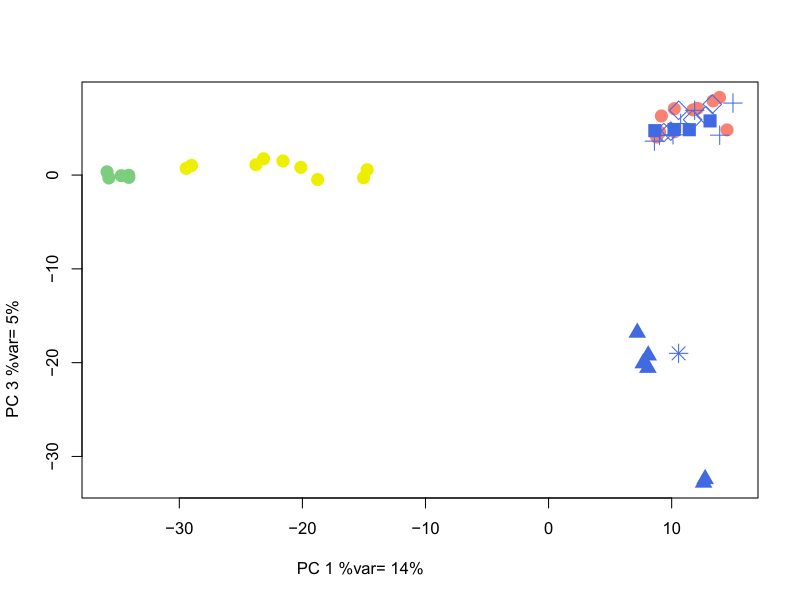

**Figure S3 (cont.) – (A) Percentage of variance explained by each principal component (PC)** on the Principal Components Analysis (PCA) (Figure 2 A and B). **(B) PC1 and PC3 of the Principal Component Analysis.**

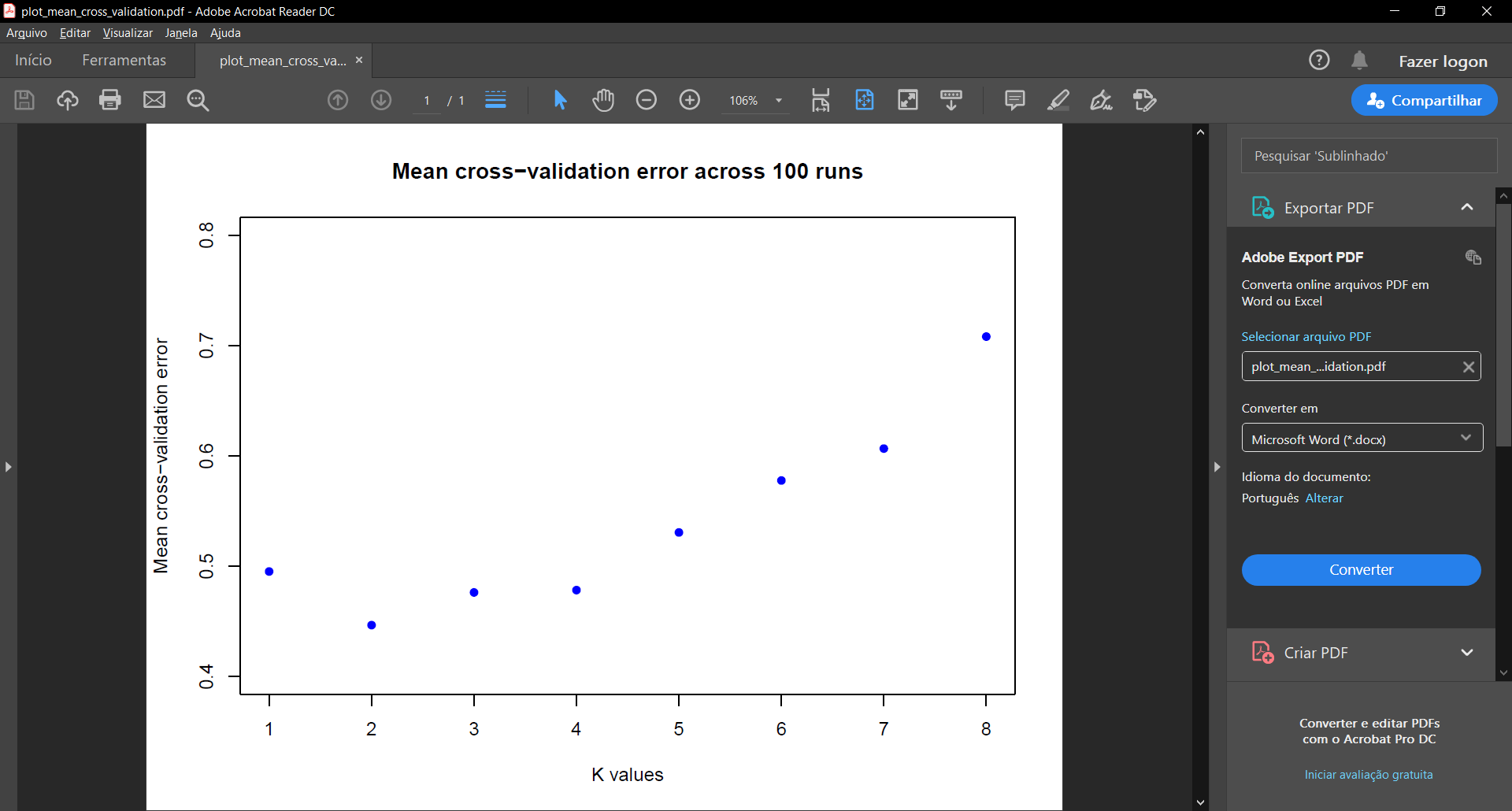

**Figure S4 – Mean cross-validation error across 100 ADMIXTURE runs.** For each value of K, the point represents the mean cross-validation error of the 100 ADMIXTURE runs performed.

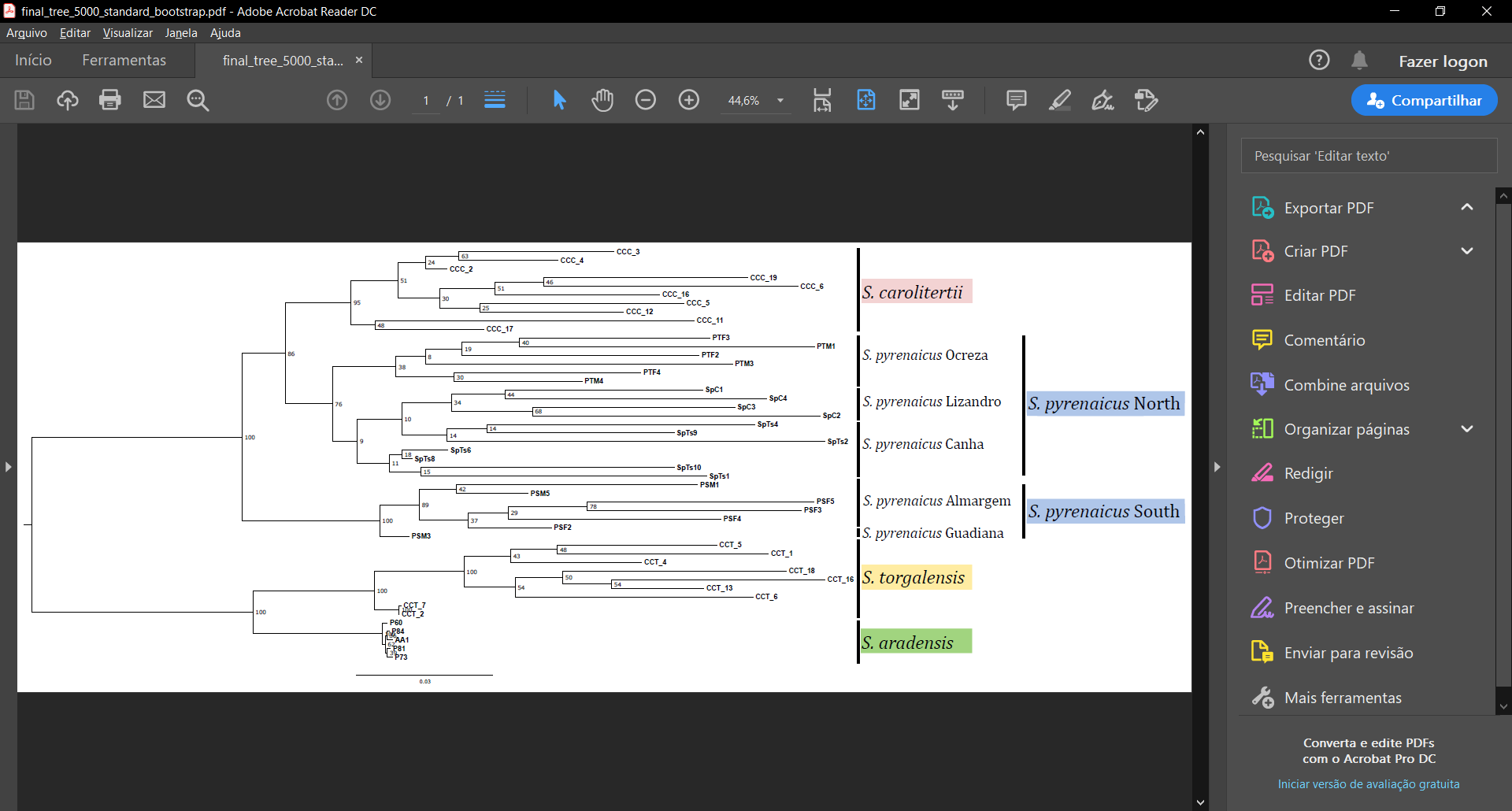

**Figure S5 – Maximum Likelihood Phylogeny.** Phylogenetic tree was obtained using IQ-TREE with 5 000 non-parametric bootstraps. Bootstrap support is included above each branch.

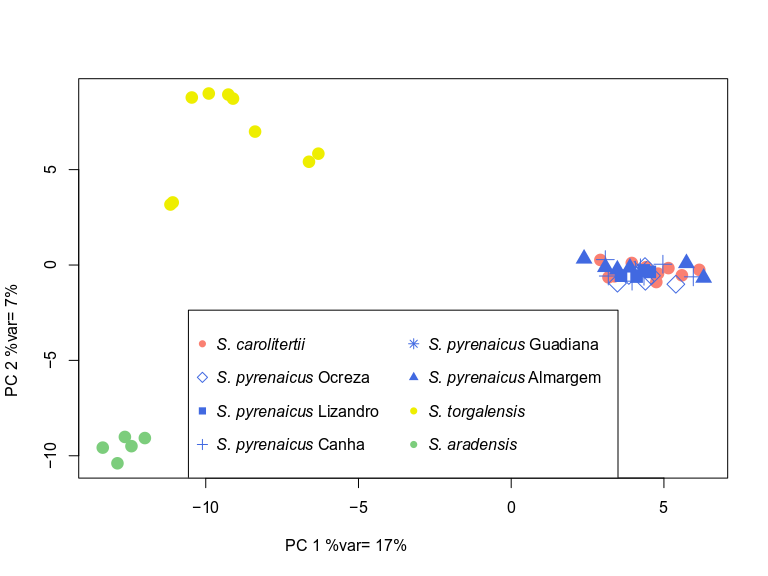

**(A)**

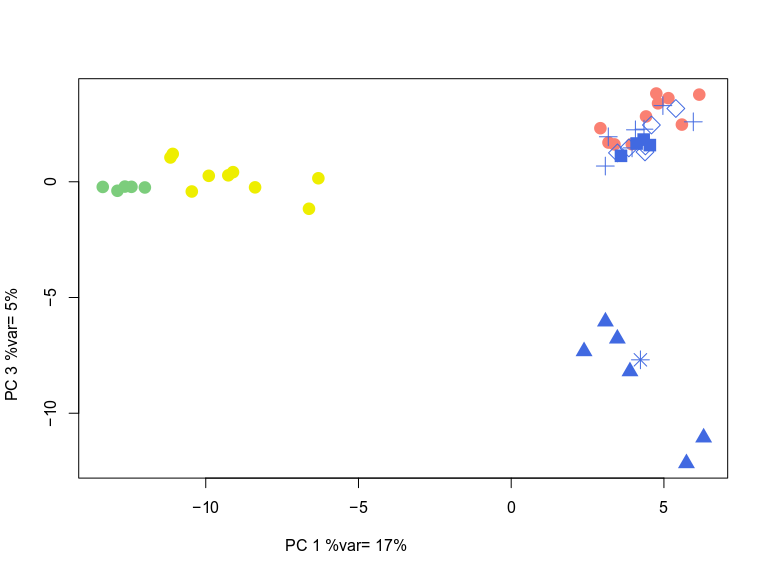

**(B)**

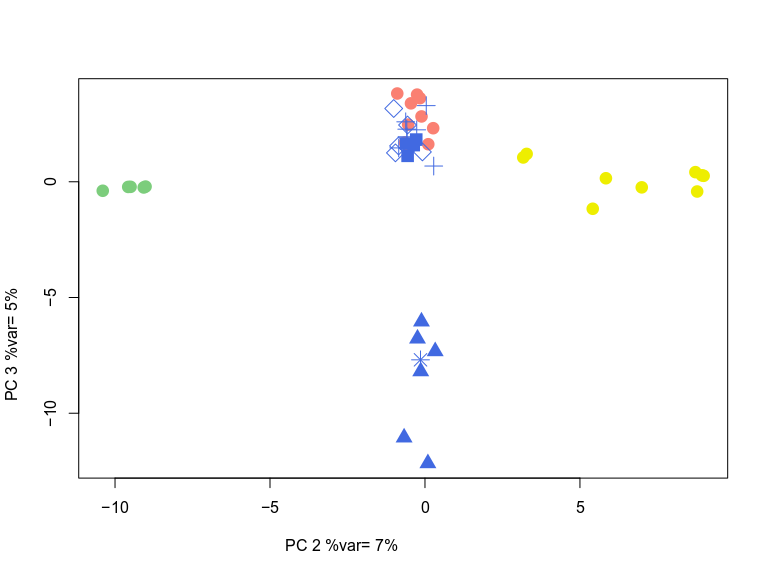

**(C)**

**Figure S6 –** **Results for the first three components of the PCA performed on the dataset with only one SNP per block: (A)** PC1 and PC2; **(B)** PC1 and PC3; **(C)** PC2 and PC3. Each point corresponds to one individual. The clusters observed were the same obtained with the larger dataset (Figure 2A and B). **(D)** **Percentage of variance explained by each principal component (PC).**

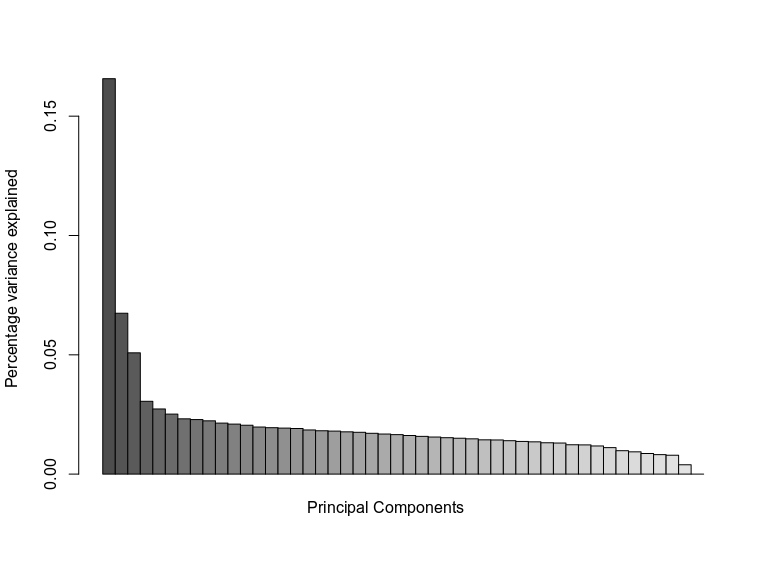

**(D)**

**Figure S6 (cont.) –** **Results for the first three components of the PCA performed on the dataset with only one SNP per block: (A)** PC1 and PC2; **(B)** PC1 and PC3; **(C)** PC2 and PC3. Each point corresponds to one individual. The clusters observed were the same obtained with the larger dataset (Figure 2A)*.* **(D)** **Percentage of variance explained by each principal component (PC)*.***

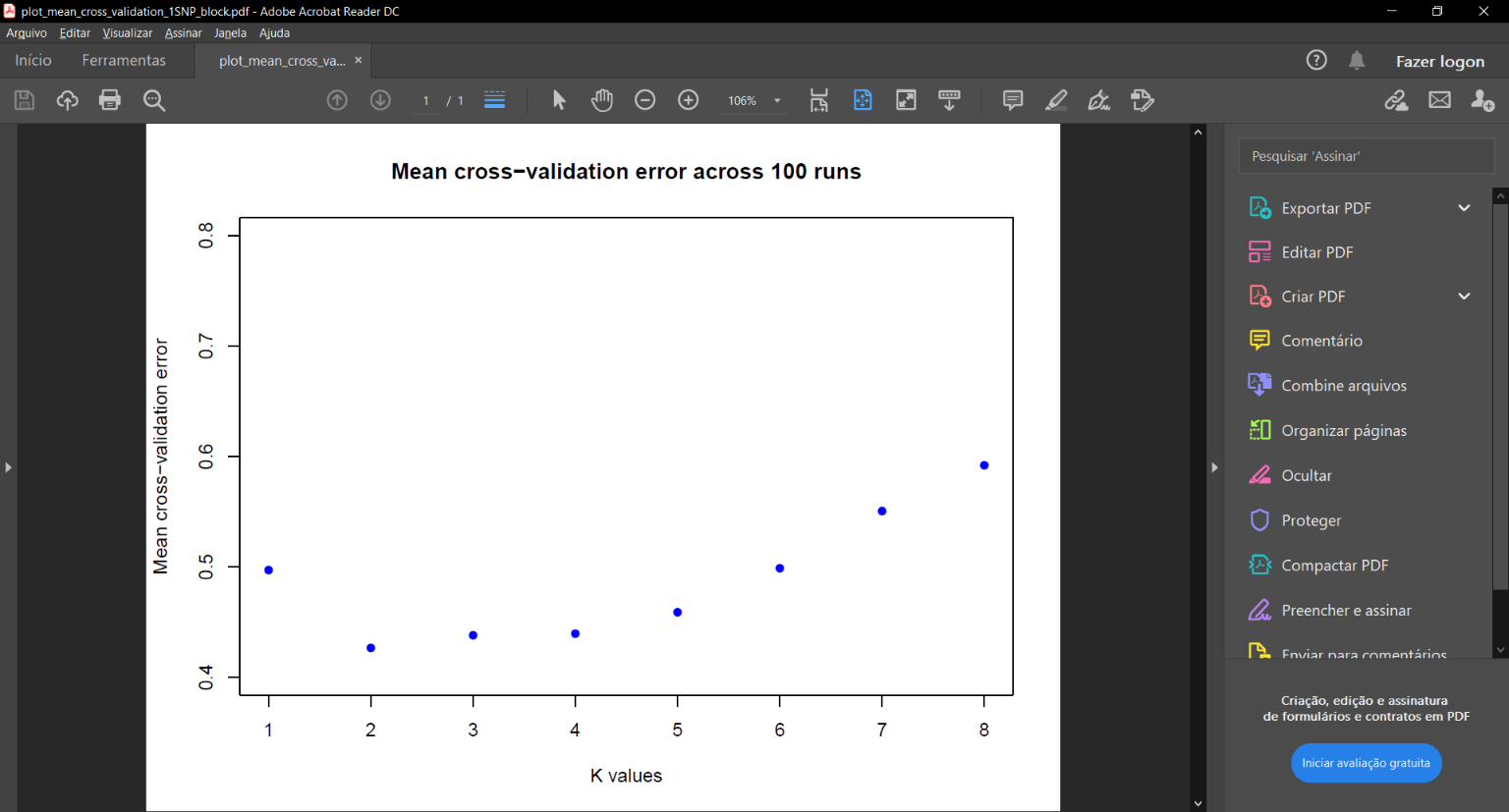

**(A)**

**(B.1)**

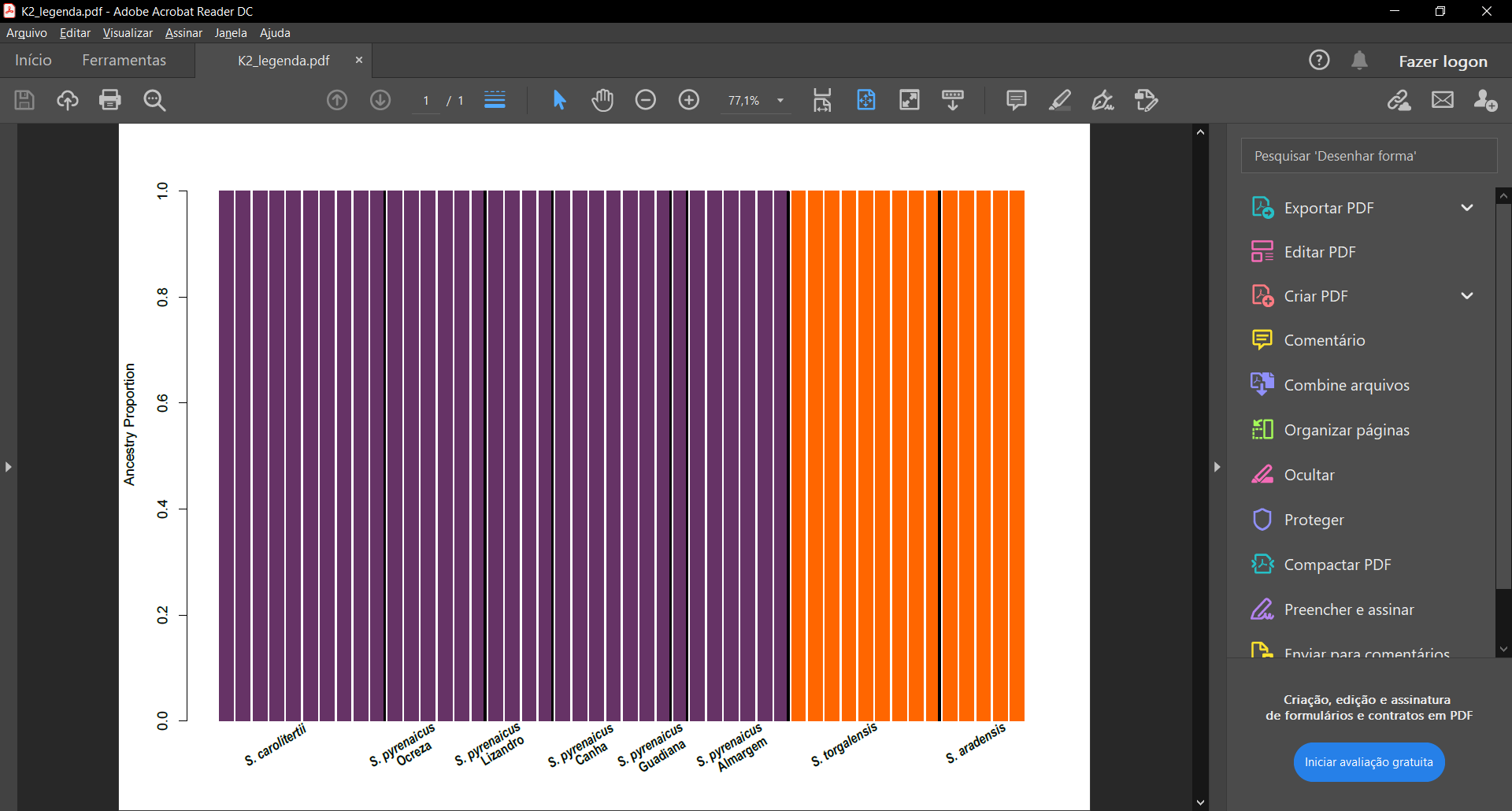

**(B.2)**

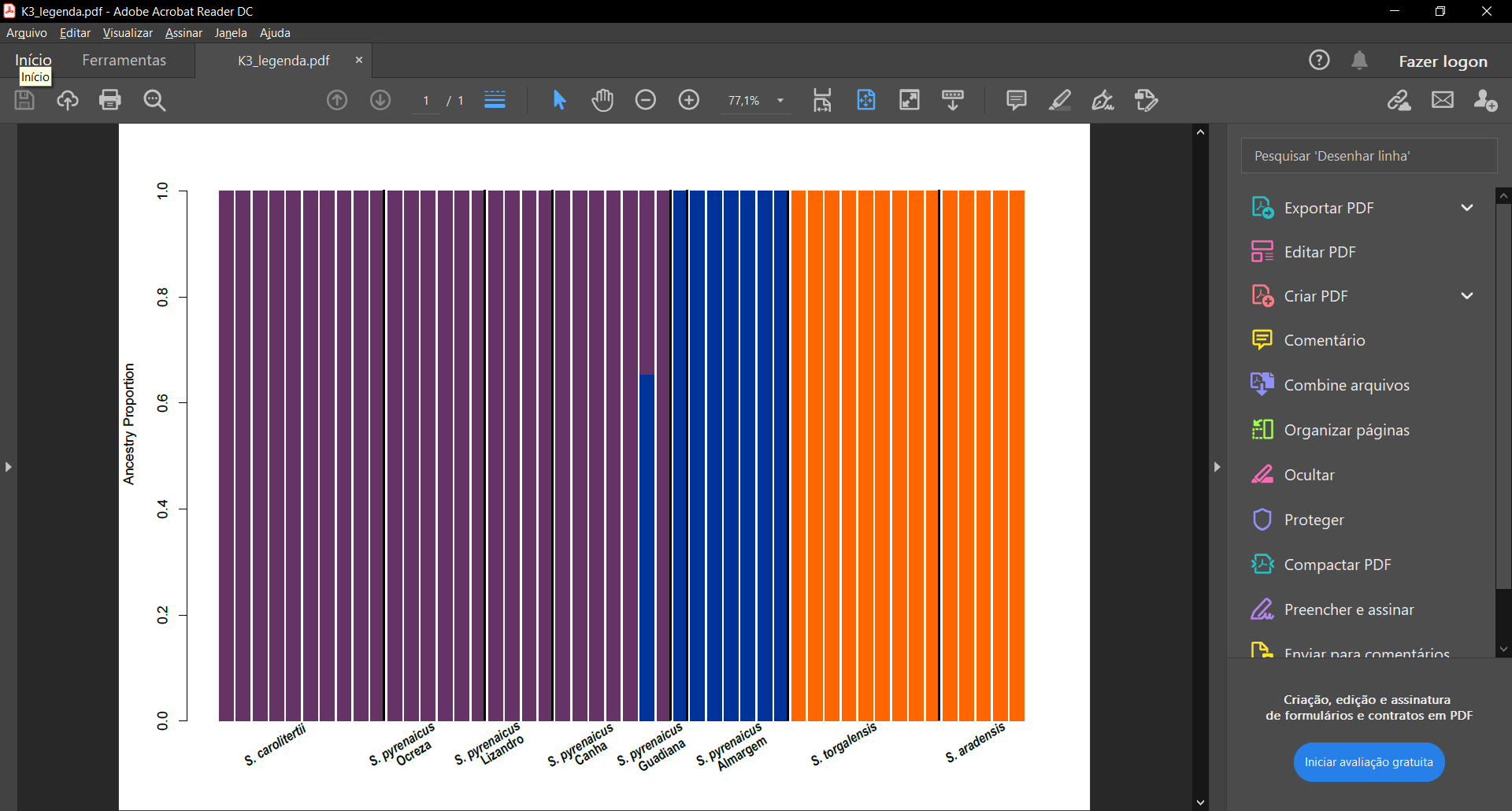

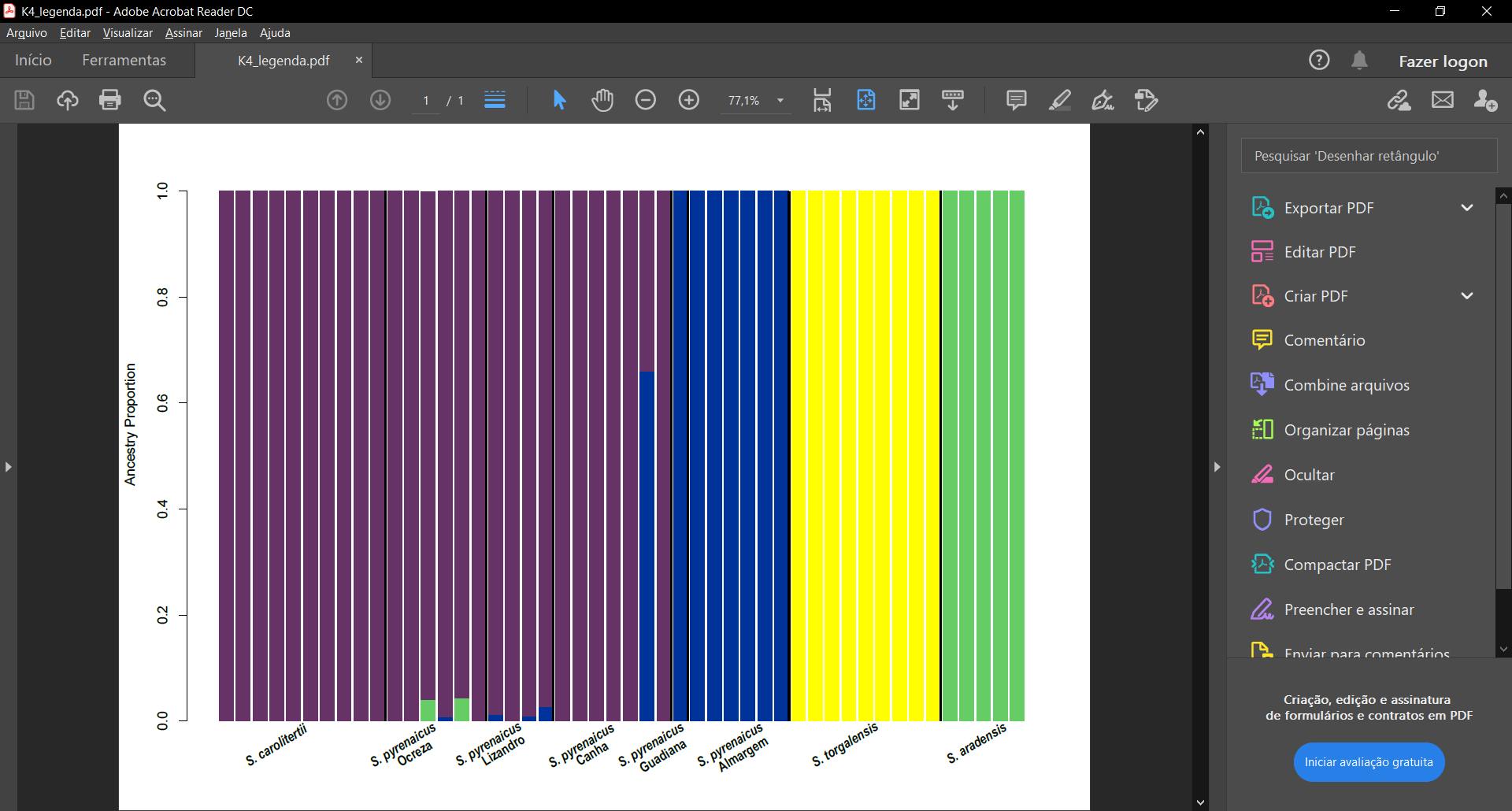

**(B.3)**

**(B.4)**

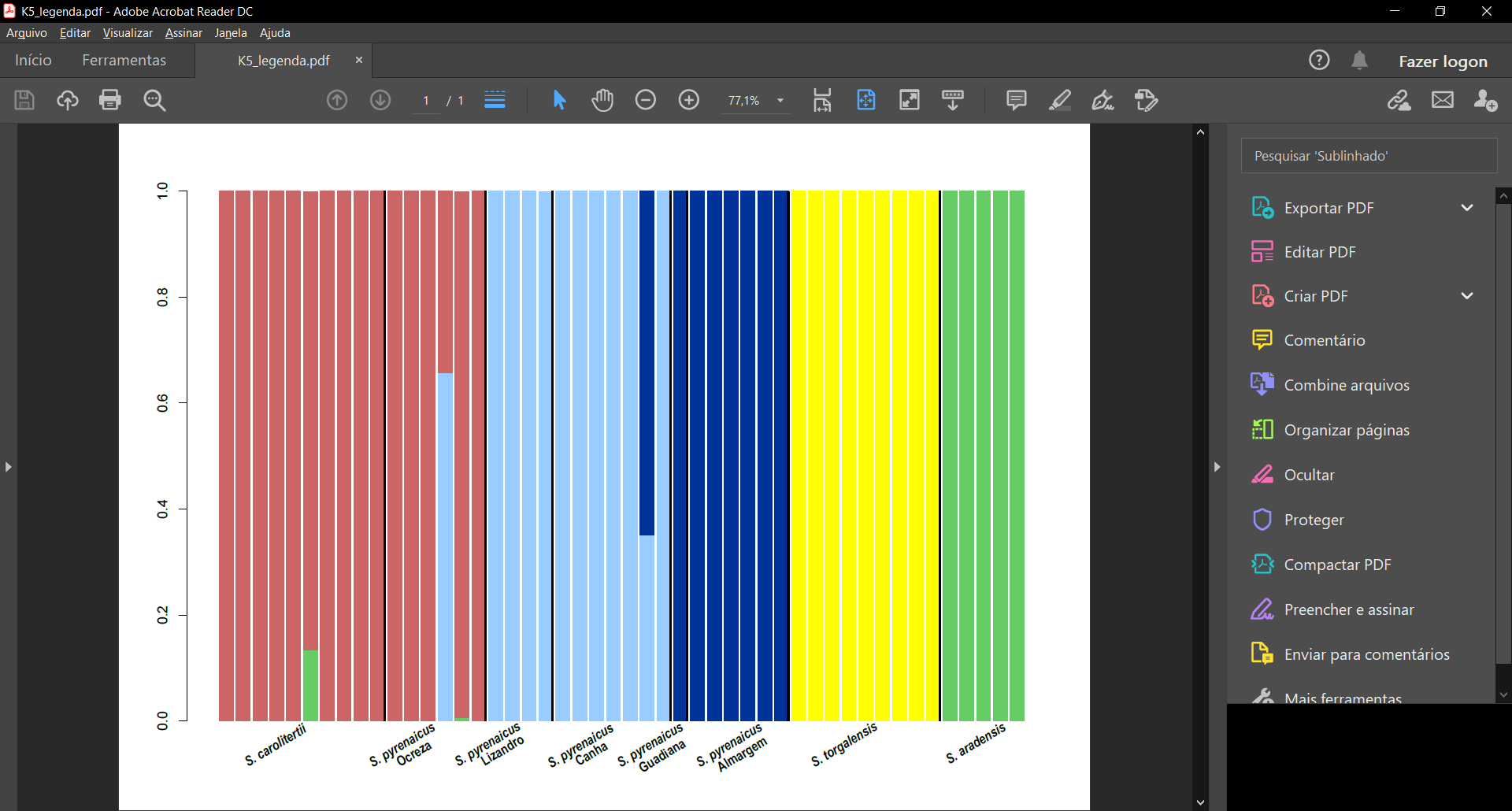

**Figure S7 – ADMIXTURE for the dataset with only one SNP per block. (A) Mean cross-validation error across 100 ADMIXTURE runs.** For each value of K, the point represents the mean cross-validation error of the 100 ADMIXTURE runs performed. **(B) Individual ancestry proportions inferred with ADMIXTURE.** **(B.1) K=2; (B.2) K=3; (B.3) K=4; (B.4) K=5.** Each vertical bar corresponds to one individual and the proportion of each colour corresponds to the estimated ancestry proportion from a given cluster. Individuals are grouped from north to south per sampling locations separated by black lines.

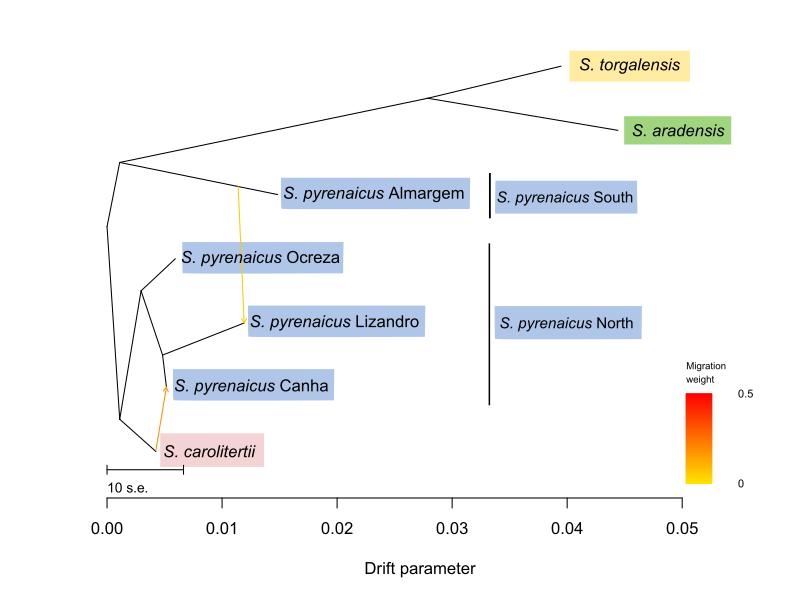

**Figure S8 - Tree graph obtained with TreeMix for the dataset with one SNP per block.** This is an unrooted tree and branch lengths are represented in units of genetic drift, i.e. the longer a given branch the stronger the genetic drift experienced in that lineage. Arrows represent migration events.

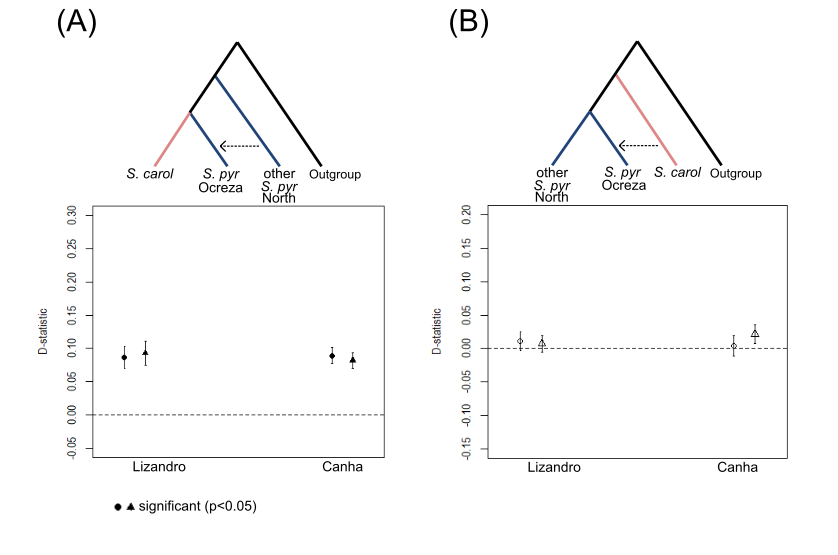

**Figure S9 - Results of the D-statistic calculated for the northern most sampling location of the northern S. pyrenaicus (Ocreza).** For each topology, the results are presented according to the other S. pyrenaicus Nroth sampling locations used (Lizandro and Canha). “S.carol” stands for S. carolitertii and “S.pyr Ocreza” stands for S. pyrenaicus Ocreza. Results obtained with each outgroup are represented by a different symbol (circles for S. torgalensis and triangles for S. aradensis). Full symbols represent significant D values (p<0.05).

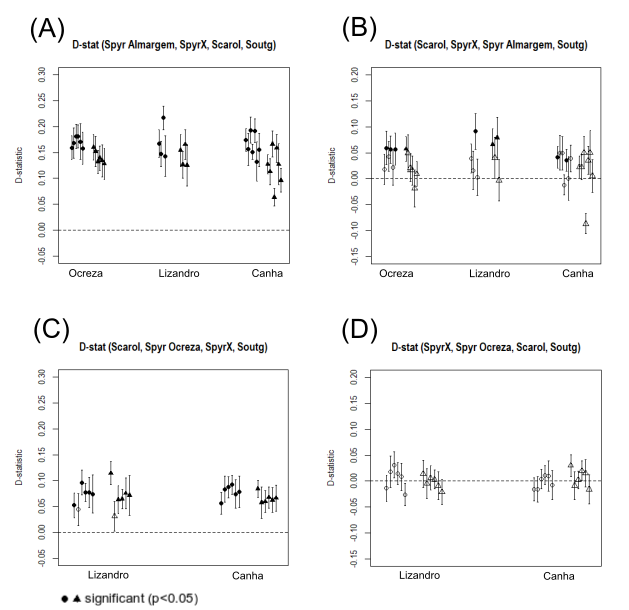

**Figure S10 -** **Results of the D-statistic calculated per individual for different topologies.** (A) and (B) correspond to topologies (A) and (B) on Figure 4. (C) and (D) correspond to topologies (A) and (B) on Figure S9. “S.carol” stands for S. carolitertii, “S.pyr Almargem” stands for S. pyrenaicus Almargem and “S.pyr Ocreza” stands for S. pyrenaicus Ocreza. Results obtained with each outgroup are represented by a different symbol (circles for S. torgalensis and triangles for S. aradensis). Full symbols represent significant D values (p<0.05).

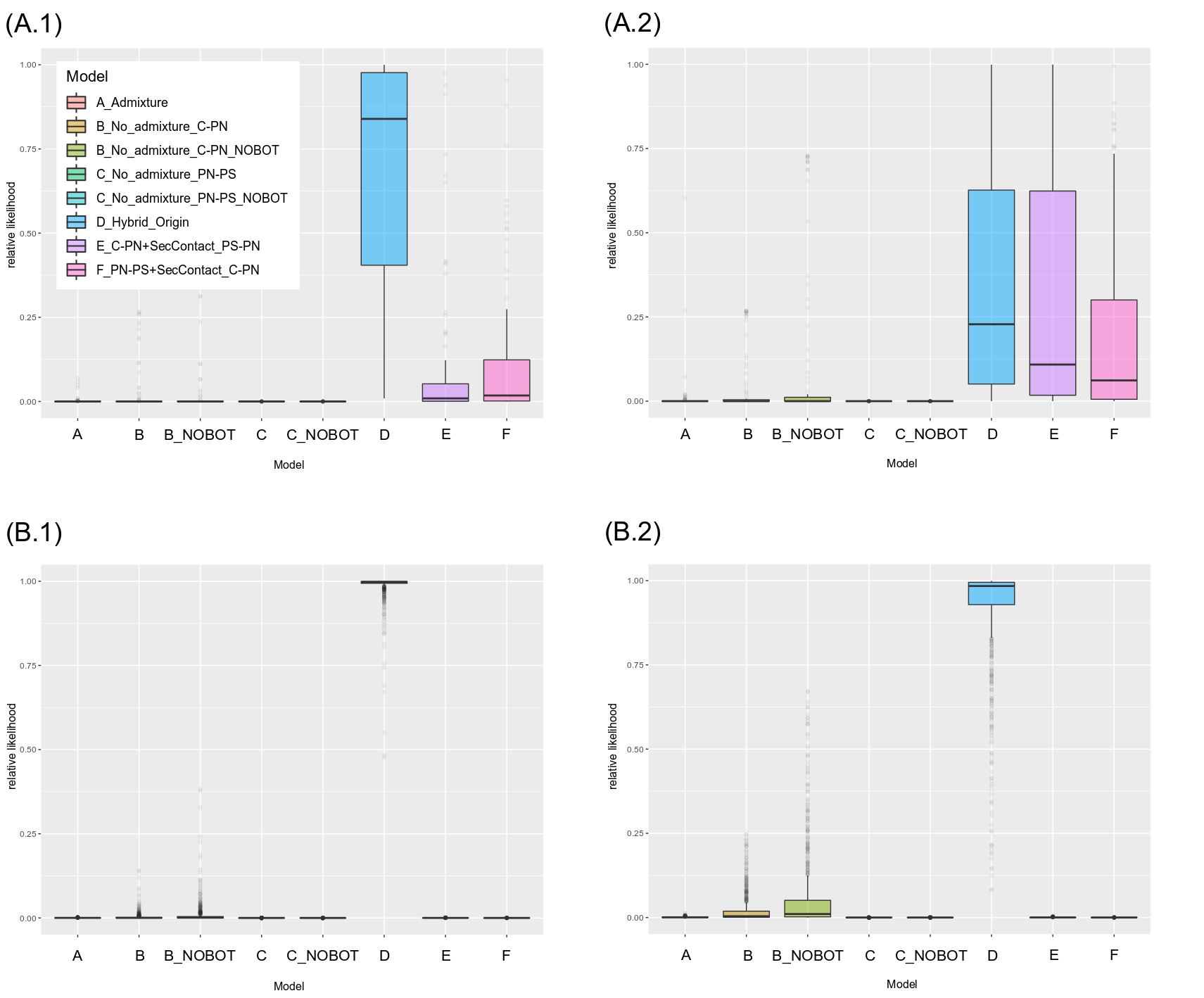

**Figure S11– Relative likelihoods of demographic models based on AIC. Comparison of 8 models, including models B and C without bottlenecks, with 1000 bootstrap replicates. (A) Boxplots of relative likelihoods based on 3D-SFS with** **all SNPs per block with n=3 S. carolitertii, n=4 S. pyrenaicus North and n=3 S. pyrenaicus South individuals.** Size of the joint 3D-SFS is SFSsize=441. Number of SNPs in observed SFS ranged from 8505 to 9135. SNPs/SFSsize = 19.29. Due to the large size of the 3D-SFS entries with less than a minimum threshold of SNPs were pooled together. Two different thresholds were considered. Boxplot of the relative likelihoods obtained with the 1000 bootstrap replicates with (A.1) minimum threshold of 1 (all entries with 1 SNP were pooled together); and (A.2) minimum threshold of 5 (all entries with 5 or less SNPs were pooled together). **(B) Boxplots of relative likelihoods based on 3D-SFS with** **1 SNP per block with n=2 S. carolitertii, n=3 S. pyrenaicus North and n=2 S. pyrenaicus South individuals.** SFS size=175. Number of SNPs in observed SFS was 2691 in all 1000 bootstrap replicates, as 1 SNP per block was sampled in all replicates. SNPs/SFSsize = 15.37. Boxplot of the relative likelihoods obtained with the 1000 bootstrap replicates with (B.1) minimum threshold of 1 (all entries with 1 SNP were pooled together); and (B.2) minimum threshold of 5 (all entries with 5 or less SNPs were pooled together). Relative likelihoods indicate that model D, consistent with a hybrid origin, reaches the highest relative likelihood irrespective of the number of SNPs per block and minimum threshold.

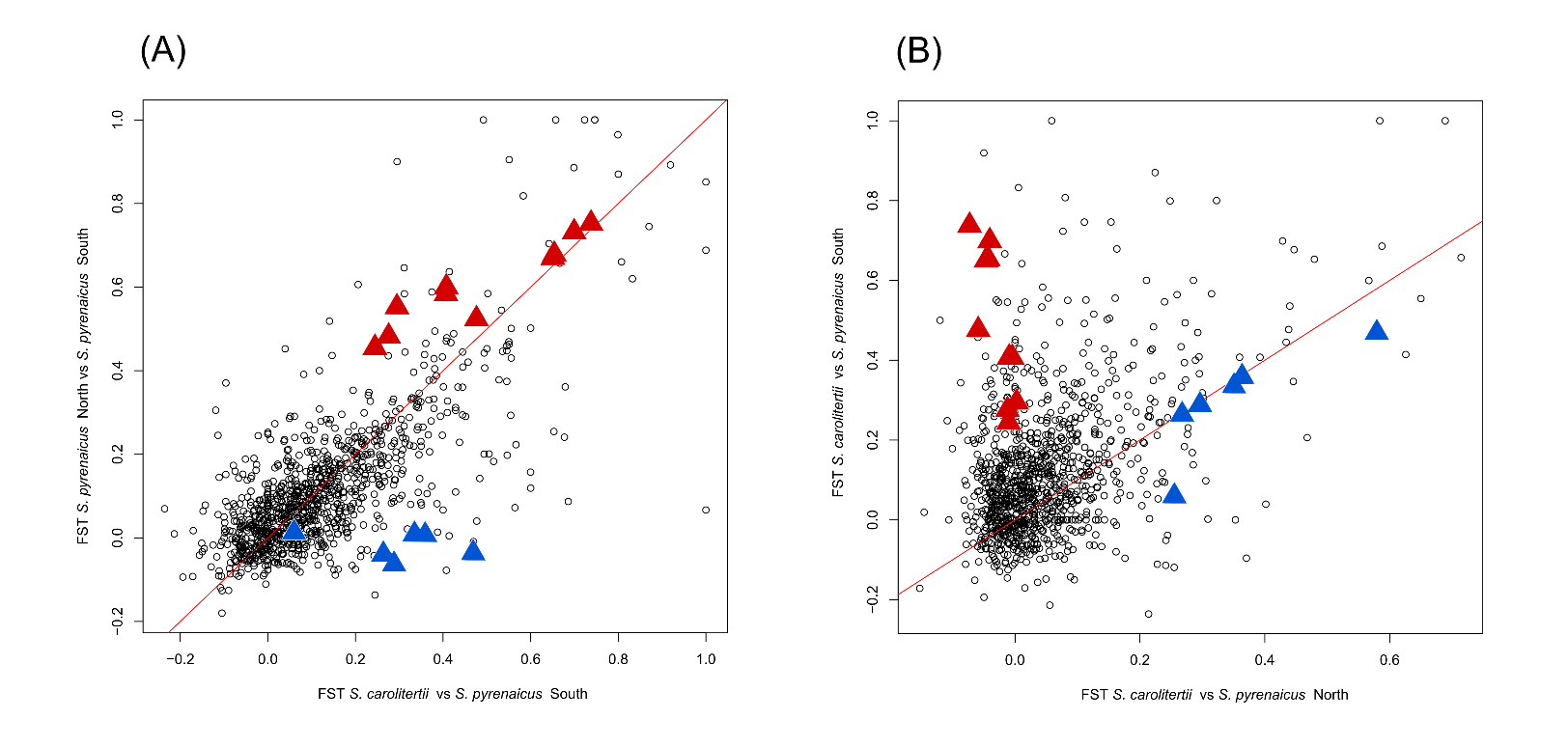

**Figure S12 –** **Outlier loci detected in *S. pyrenaicus* North.** Red triangles represent loci from *S. carolitertii* selected for in *S. pyrenaicus* North. Blue triangles represent loci from *S. pyrenaicus* South selected for in *S. pyrenaicus* North.
